# Redirecting cytotoxic lymphocytes to breast cancer tumors via metabolite-sensing receptors

**DOI:** 10.1101/2025.03.21.644686

**Authors:** Young-Min Kim, Reece V. Akana, Chang Sun, Olivia Laveroni, Livnat Jerby

## Abstract

Insufficient infiltration of cytotoxic lymphocytes to solid tumors limits the efficacy of immunotherapies and cell therapies. Here, we report a programmable mechanism to mobilize Natural Killer (NK) and T cells to breast cancer tumors by engineering these cells to express orphan and metabolite-sensing G protein-coupled receptors (GPCRs). First, in vivo and in vitro CRISPR activation screens in NK-92 cells identified *GPR183*, *GPR84*, *GPR34*, *GPR18*, *FPR3*, and *LPAR2* as top enhancers of both tumor infiltration and chemotaxis to breast cancer. These genes equip NK and T cells with the ability to sense and migrate to chemoattracting metabolites such as 7α,25-dihydroxycholesterol and other factors released from breast cancer. Based on Perturb-seq and functional investigations, GPR183 also enhances effector functions, such that engineering NK and CAR NK cells to express GPR183 enhances their ability to migrate to, infiltrate, and control breast cancer tumors. Our study uncovered metabolite-based tumor immune recruitment mechanisms, opening avenues for spatially targeted cell therapies.

## INTRODUCTION

Mapping the logic of immune cell migration and identifying mechanisms to redirect immune cells to specific sites in the body and within a target tissue is critical, as the location of immune cells dictates whether and how these cells elicit their effect, impacting the course of the entire immune response. Demonstrating this, while immunotherapies and cell therapies have transformed the therapeutic landscape in several cancer types (1–3), insufficient recruitment and infiltration of cytotoxic lymphocytes to the tumor is still limiting the efficacy of these treatments in solid tumors in many patients (4–7), underscoring the need and potential of “cell delivery” solutions.

Immune cell migration and infiltration rely on chemical, mechanical, and electrical signals and the capacity of the immune cells to sense, process, and respond to these signals (8–10). The intrinsic differences between different immune cell types and states result in differential responses to such signals and different spatial distributions at the whole-body and tissue levels and within solid tumors. The main genes thought to orchestrate immune cell recruitment to solid tumors are chemokines and their cognate receptors expressed by immune cells, as well as genes encoding for adhesion proteins as selectins and integrins that mediate transmigration and infiltration (11,12). Yet, while a lot is already known about cell migration and infiltration, our ability to mobilize immune cells to specific sites and into solid tumors is still limited.

Forward genetic screens offer a powerful framework to optimize different cell functions of interest via directed evolution, such that genetic variation is introduced (e.g., via CRISPR or other tools) and the pool of engineered cells undergoes selection to identify the cells that are optimal at performing a “task” of interest. In this context, gain-of-function studies are equally important to loss-of-function ones because expressing genes in a cellular context where they are not endogenously (over)expressed can reprogram cells in ways that cannot be achieved based on gene inhibition or knockout, generate new mechanisms, or hijack those that are active in different cellular contexts. While a variety of CRISPR knockout (13–15), interference, and activation (16), as well as ORF (open reading frame) (17) and shRNA screens, have been performed to study different immune cell functions, to date, there has not been any gain-of-function screen to identify genes that can enhance immune cell infiltration to solid tumors.

Here, we performed an array of in vivo and in vitro CRISPR activation screens in NK-92 cells and identified a programmable mechanism to recruit cytotoxic lymphocytes [including both Natural Killer (NK) and CD8 T cells] to breast cancer tumors based on the activation of G protein-coupled receptors (GPCRs) that sense chemoattracting metabolites. We show that top hits equip NK-92 cells, primary NK cells, and primary CD8 T cells with the ability to sense and migrate toward chemoattracting metabolites and factors released from breast cancer cells and tumors. Via Perturb-seq screening and functional investigations, we show that *GPR183* enhances NK cell effector functions, such that engineering NK and chimeric antigen receptor (CAR) NK cells to express *GPR183* enhances their ability to migrate to, infiltrate, and control breast cancer tumors in mice.

Thus, while aberrant metabolism has been a well-known hallmark of cancer for many decades now (19–22), our study shows that it is possible to leverage cancer aberrant metabolism to recruit cytotoxic lymphocytes to the tumor via chemoattracting metabolites, opening new avenues for the design and development of spatially targeted cell therapies.

## RESULTS

### CRISPR activation screens in NK-92 cells identified metabolite-sensing GPCRs as enhancers of cell migration to breast cancer tumors

To identify mechanisms to mobilize NK cells to solid tumors, we conducted in vivo CRISPR activation screens designed to identify genes that significantly enhance NK cell ability to migrate to and populate breast cancer tumors in mice (**Figure 1a**). We conducted the screens in a clinically applicable human NK cell line (23,24) (NK-92) that has been widely used for CAR NK studies and in clinical trials (25,26). To power the screens, we first identified genes that are more likely to drive recruitment to breast cancer tumors in patients by analyzing scRNA-seq data collected from 22 breast cancer patients with matched samples from tumor and peripheral blood (27). For each immune cell type with sufficient representation in the data, we identified genes that were significantly overexpressed in the tumor compared to blood samples [Benjamini-Hochberg (BH) False Discovery Rate (FDR) < 0.05, log fold change > 0.1, MAST test (28); **Figure 1b** and **Supplementary Figure 1a**]. We did not constrain these analyses only to NK cells because we hypothesized that there may be genes that facilitate or drive the recruitment of other (non-NK) immune cells to solid tumors and can elicit similar effects in NK cells if expressed exogenously [e.g., via CRISPR activation or an Open Reading Frame (ORF)].

**Figure 1.**
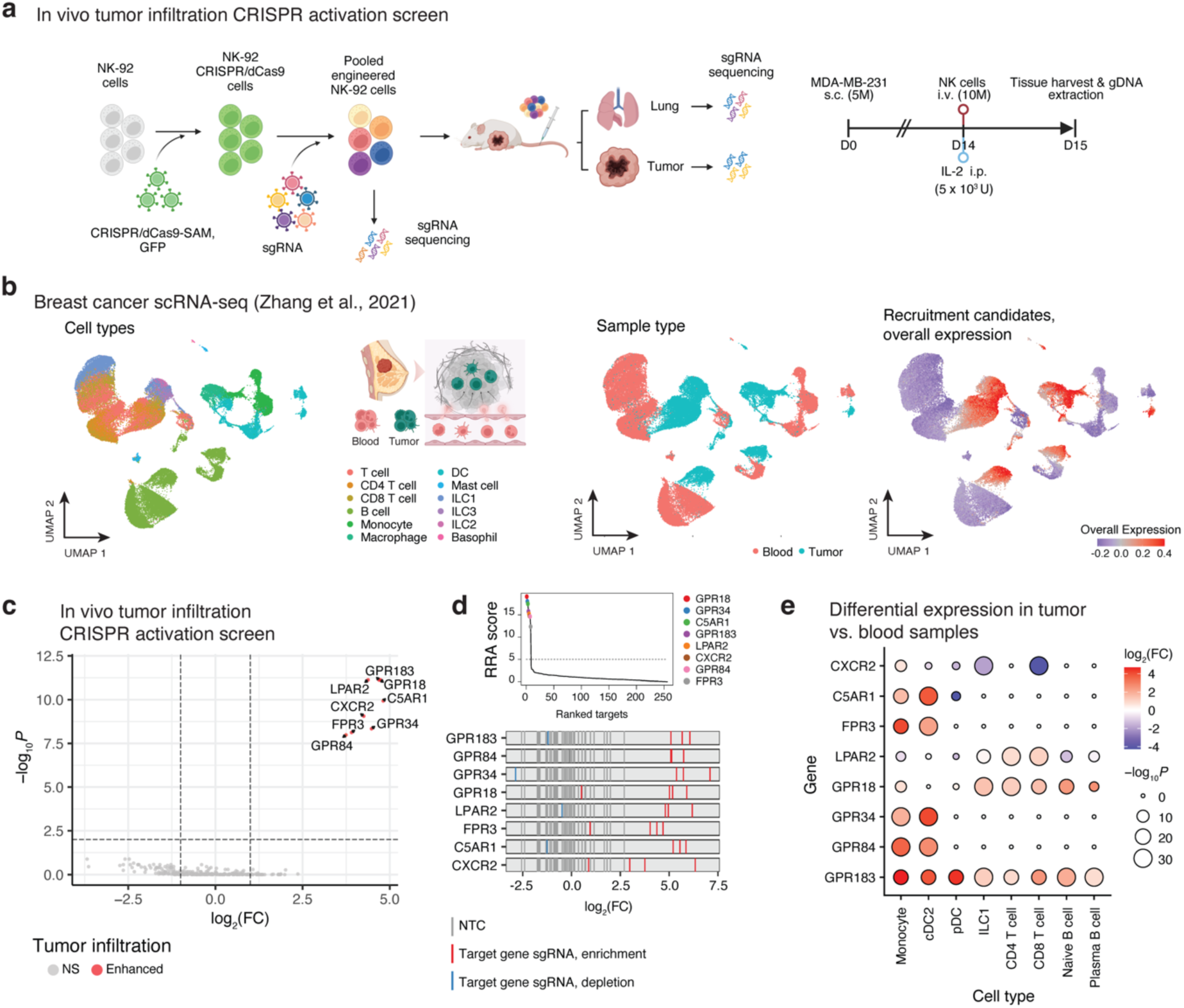
In vivo CRISPR activation screens in NK-92 cells identified metabolite-sensing GPCRs as enhancers of NK cell infiltration to breast cancer tumors. **a,** Experimental scheme of in vivo CRISPR activation screen. **b,** Uniform Manifold Approximation and Projection (UMAP) of the scRNA-seq data from breast cancer patients that was analyzed to identify candidate tumor infiltration enhancers. Each dot corresponds to a cell, colored by cell type (left), sample type (middle), and the overall expression of the genes identified as overexpressed in the immune cells residing in the tumor compared to the blood samples (right; **Methods**). ILC: innate lymphoid cell. **c,** CRISPR activation tumor infiltration screen results are shown as the significance (y-axis) and log-transformed fold-change (x-axis) of each target gene (dot) in the tumor compared to lung samples based on MAGeCK (36,37). Target genes whose sgRNAs were significantly or not significantly (NS) enriched in the tumor compared to lung samples are colored in red and grey, respectively. There were no target genes showing a significant depletion in the tumor compared to lung samples, aligned with the screen being a strong positive selection screen. **d,** Top: robust rank aggregation (RRA) scores of all target genes matching the analysis shown in **(c)**; bottom: log-transformed fold change of sgRNAs in tumor versus lung samples shown for sgRNAs targeting indicated genes. **e,** Differential expression of top hits in tumor versus blood samples in breast cancer patients based on scRNA-Seq data shown in **(b)**. Differential expression analyses were performed using the MAST test (28) (**Methods**). The graphics in **(a)** and **(b)** were created with Biorender.com.

To test the candidate recruitment enhancing genes, we generated a library of 1,070 CRISPR activation single guide RNAs (sgRNAs) targeting a total of 256 genes [4 sgRNAs per target gene, and 46 non-targeting control (NTC) sgRNAs], including both candidate genes identified in patients as well as additional genes with well-established roles in NK biology or immune chemoattraction and invasion, as proteases, adhesion molecules, integrins, and chemokine receptors (**Supplementary Figure 1a** and **Supplementary Table 1**). We transduced NK-92 cells to stably express CRISPR/dCas9-SAM (synergistic activation mediator) system (29) and confirmed sgRNA-based gene overexpression at the protein level via flow cytometry (**Supplementary Figure 1b**). We then transduced the CRISPR/dCas9-SAM NK-92 cells with the sgRNA library at a low multiplicity of infection (MOI < 0.3) to generate a pool of engineered NK cells, each carrying a single sgRNA to activate one of the 256 genes (**Supplementary Table 1**).

Intravenously transferring the pool of engineered NK cells to breast cancer tumor-bearing mice, we performed in vivo screens to identify engineered NK cells with superior ability to migrate to and infiltrate the tumors compared to control NK cells (transduced with NTC sgRNAs). We performed the screens in non-obese diabetic scid gamma (NSG) female mice engrafted with triple-negative breast cancer cells (MDA-MB-231), either subcutaneously or orthotopically in the left inguinal (4^th^) mammary fat pad and in transgenic NOG mice that express the human IL2 gene (hIL2-NOG) and are better at retaining NK cells. In all three models, we collected tumor and lung samples 24 hours after the NK cell adoptive transfer for genomic DNA (gDNA) extraction and sgRNA sequencing. As control samples, we sequenced both the NK cell pool from which samples were drawn for in vivo transfer (denoted as the “pre-injection pool”), as well as the lung tissue samples. We chose the lung as a control site because it has been reported to accumulate and sequester adoptively transferred tumor-reactive lymphocytes, both decreasing tumor control (30–32) and resulting in severe lung toxicities (33–35). All samples were processed and sequenced together and demultiplexed post-sequencing (**Figure 1a**).

We performed animal-matched paired statistical data analyses to identify gene activations that resulted in substantial, significant, and reproducible tumor enrichment, both compared to the lung and compared to the pre-injection NK cell pool (using MAGeCK (36,37), **Methods**). Eight genes – *GPR183*, *GPR84*, *GPR34*, *GPR18*, *LPAR2*, *FPR3*, *C5AR1*, and *CXCR2* – were consistently ranked as the top hits (BH FDR < 1*10^-7^, Fisher test) in all three screens (**Figure 1c,d, Supplementary Figure 2**, and **Supplementary Table 2**). All the hits were initially identified as candidates based on our scRNA-seq data (27) analyses, as they were significantly overexpressed in breast cancer tumors compared to blood samples in the cell types where they are endogenously expressed. Based on this data *C5AR1*, *FPR3*, *GPR34*, and *GPR84* are primarily expressed by myeloid and dendritic cells, *GPR18* is mostly expressed by lymphocytes [e.g., T cells, B cells, and innate lymphoid cells (ILCs)], *CXCR2* and *LPAR2* show a relatively low expression across all immune cell types in this dataset, and *GPR183* is expressed across a broader set of immune cell types (**Figure 1e** and **Supplementary Figure 3**).

Collectively, the screens repeatedly identified eight GPCRs as enhancers of NK-92 cell migration to breast cancer tumors in mice, with only one of them being a chemokine receptor. This prompts us to investigate the molecular mechanisms that mediate these effects.

### Top hits redirect NK cells to migrate to chemoattracting metabolites and factors released from breast cancer cells

According to the International Union of Basic and Clinical Pharmacology (IUPHAR) (38) classification, all eight hits identified in our screens are GPCRs coupled with G_i/o_ protein, a primary signal transducer for most conventional chemokine receptors. Yet, unlike chemokine receptors, five out of the eight hits—*GPR183*, *GPR84*, *GPR34*, *GPR18*, and *LPAR2—*have been reported to bind and respond to bioactive metabolites, including lipid, fatty acid, and cholesterol derivatives (39–42).

Focusing on *GPR183*, *GPR84*, and *GPR34*, we examined whether these GPCRs sense bioactive metabolites and redirect NK cell migration to these metabolites. First, using the PRESTO-tango system (43), we generated reporter lines for these three hits as well as *C5AR1* and *CXCR2* (top hits marking well-established complement- and chemokine-based chemotaxis that we used here as controls). Using the PRESTO-tango reporter lines, where the conformational change of the GPCR in response to its agonist ligand is converted to a bioluminescence signal (**Figure 2a**), we confirmed that 7α,25-dihydroxycholesterol (7α,25-OHC), lysophosphatidylserine (LysoPS), 6- OAU, complement component 5a (C5a), and interleukin 8 (IL8) are the agonists of *GPR183*, *GPR34*, *GPR84*, *C5AR1,* and *CXCR2*, respectively (**Figure 2b**). We then generated NK-92 lines that constitutively express each one of these hits via CRISPR activation or ORF (**Supplementary Figure 4**). Using these syngeneic NK-92 lines, we tracked and quantified migration to the different agonists, showing *GPR183-*, *GPR34-*, *GPR84-*, *C5AR1-,* and *CXCR2*- dependent migration to 7α,25-OHC, LysoPS, medium 6-OAU, C5a, and IL8, respectively (**Figure 2c**).

**Figure 2.**
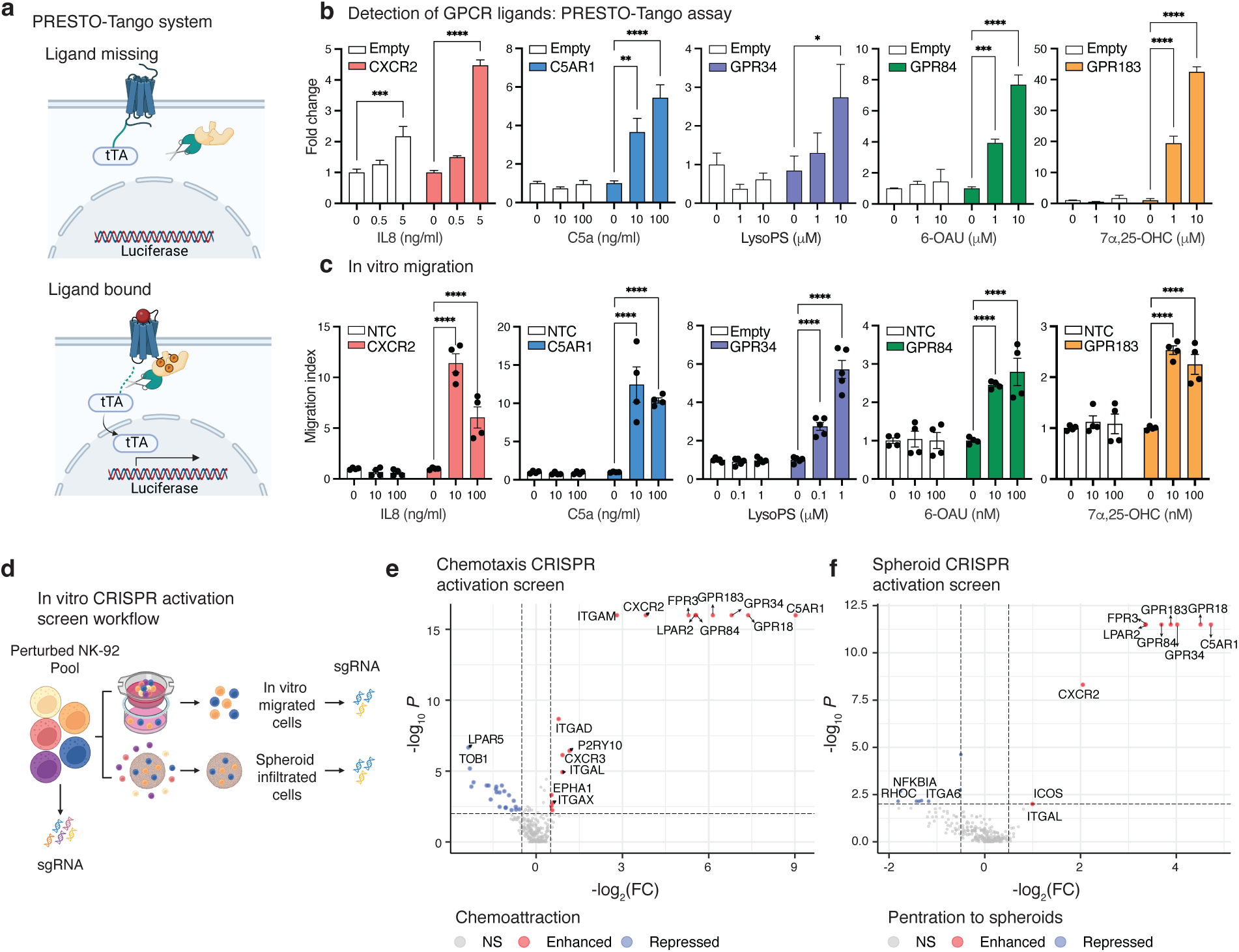
Tumor infiltration hits enhance NK cell sensing of and migration to chemoattracting metabolites and factors released from breast cancer cells. **a,** Schematics of PRESTO-tango reporter cell lines where GPCR conformational changes in response to a ligand result in luciferase transcription for bioluminescence readouts. **b,** PRESTO-tango reporter cell lines generated for CXCR2, C5AR1, GPR34, GPR84, and GPR183 confirm IL8, C5a, LysoPS, 6- OAU, and 7α,25-OHC as the respective GPCR agonists (n = 3). **c,** GPCR-driven migration of NK- 92 cells to the respective ligand (n = 4 or 5). Data are representative of two independent experiments and presented as the mean ± s.e.m. One-way analysis of variance (ANOVA) was performed with Dunnett’s multiple comparisons for (**b**) and (**c**) (\*\*\*\**P* < 0.0001, \*\*\**P* < 0.001, \*\**P* < 0.01, and \**P* < 0.05). **(d-f)** Chemotaxis and spheroid infiltration CRISPR activation screens in NK-92 cells: **d,** experimental scheme, **(e,f)** screen results shown as the significance (y-axis) and fold change (x-axis) of each target gene (dot). Positive and negative values denote enrichment and depletion, respectively, in the NK cell population that migrated to the breast cancer supernatant (**e**) and infiltrated into the breast cancer spheroids (**f**). NS: Not significant. The graphics in (**a**) and (**d**) were created with Biorender.com.

Thus, we hypothesized that the hits identified may surface a different form of immune cell recruitment to the tumor that is based on chemoattracting metabolites released from the cancer cells. To test this hypothesis and systematically identify genes that enhance NK cell chemoattraction to breast cancer cells, we performed two additional in vitro screens (**Figure 2d**). The first in vitro screen was designed to identify perturbations that enhance chemotaxis to factors released from breast cancer cells (MCF7 and MDA-MB-231) using a transwell device (**Figure 2d,e**). The second in vitro screen was conducted by coculturing the pool of genetically modified NK cells with breast cancer (MCF7) spheroids and selecting for the NK cells that both migrated and penetrated to the spheroids within 6, 24, and 48 hours (**Figure 2d,f**, and **Supplementary Figure 5a**). All eight hits identified in the in vivo screen were among the top 10 hits in both the chemotaxis and spheroid in vitro screens (**Figure 2e,f, Supplementary Figure 5b-d**, and **Supplementary Table 3**). The chemotaxis screen identified several other hits, including the chemokine receptor *CXCR3,* the integrins *ITGAD*, *ITGAL*, *ITGAX,* and *ITGAM*, which are known for their role in lymphocyte adhesion and transmigration across blood vessels, and *EPHA1*, which is known for its role in NK cell migration (44). The only other hits identified in the spheroid screen were *ICOS* and *ITGAL*.

Collectively, these findings demonstrate that the GPCRs identified as enhancers of NK cell migration to breast cancer tumors in mice enhance NK cell ability to sense and migrate to chemoattracting metabolites and soluble factors released by breast cancer cells.

### GPR183 alters the NK cell transcriptome and effector functions

Tumor metabolism has often been reported to be immunosuppressive, with a recent study demonstrating that LysoPS prevents type 1 ILCs (ILC1s) activation via GPR34 (45). Thus, while our screens selected for perturbations that enhance NK cell migration to breast cancer tumors, there is no guarantee that these perturbations will not interfere with NK cell effector functions, surfacing the need to determine the impact of the hits on NK cell state and functionality. As GPCR signaling results in transcriptional regulation, we turned to examine if and how the hits impact the NK cell transcriptome by conducting a Perturb-seq (46,47) screen, where we combined scRNA-seq with matched sgRNA detection at the single cell level in NK-92 cells (**Figure 3a** and **Supplementary Figure 6a**). The Perturb-seq data confirmed on-target gene activation and demonstrated that *GPR183*, *C5AR1*, and *CXCR2* CRISPR activation result in a substantial impact on the NK-92 cell transcriptome, with hundreds of differentially expressed genes in comparison to the control NK-92 cells (transduced with NTC sgRNAs; **Figure 3b,c**, **Supplementary Figure 6b,** and **Supplementary Table 4**).

**Figure 3.**
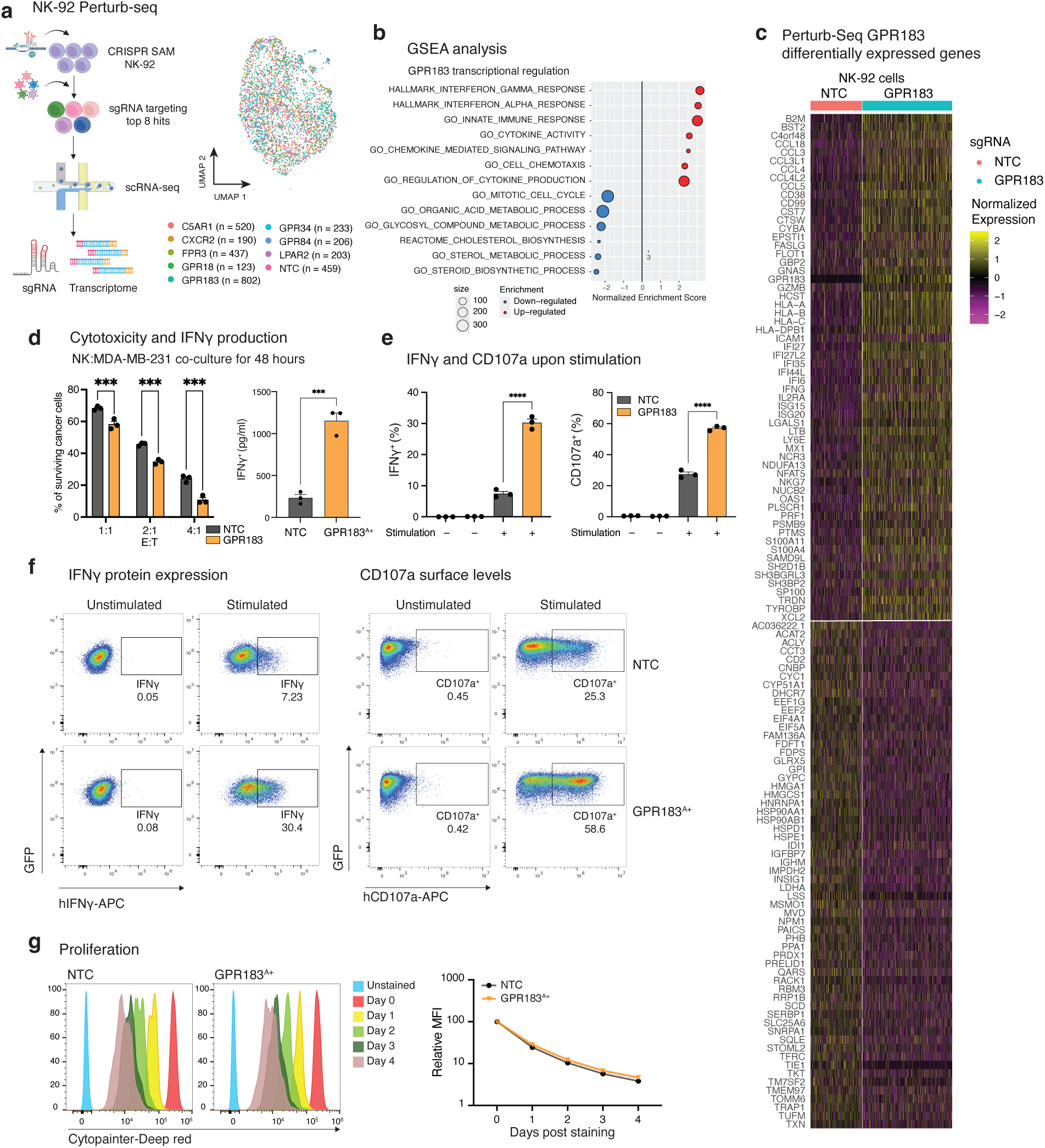
GPR183 activation alters NK cell transcriptome and improves effector functions. (**a-c**) NK-92 Perturb-seq screen of top hits. **a**, Schematics of perturb-seq workflow and UMAP of Perturb-seq data, where each dot corresponds to an NK-92 cell colored by the sgRNA target gene. **b**, Gene set enrichment analysis of genes significantly differentially expressed in GPR183^A+^ vs. NK-92 cells with NTC sgRNAs (i.e., control NK cells). **c**, Heatmap of GPR183 differentially expressed genes in NK-92 cells: Each column corresponds to an NK-92 cell with either GPR183 or NTC sgRNAs, as indicated by the topmost horizontal color bar; each row corresponds to the topmost differentially expressed genes in the GPR183^A+^ vs. controls NK-92 cells. **d**, Cytotoxicity and IFNγ secretion in GPR183^A+^ cells vs. control NK-92 cells (n = 3). Cytotoxicity to MDA-MB-231 cells was evaluated after 48 hours of coculture with the NK-92 cells at varying effector-to-target (E:T). IFNγ secretion was evaluated at 1:1 E:T. **e**, Bar plots depict flow cytometry measures of IFNγ protein expression and CD107a levels on the NK-92 cell surface obtained at baseline and upon stimulation across biological replicates (n = 3). Data are representative of two independent experiments and presented as the mean ± s.e.m. Two-tailed unpaired Students’ t-test for (**d**) and (**e**) (\*\*\*\**P* < 0.0001, \*\*\**P* < 0.001, and \*\**P* < 0.01). **f**, Flow cytometry measures of IFNγ protein expression and CD107a levels on the NK-92 cell surface in GPR183^A+^ vs. control NK cells at baseline and upon stimulation with PMA/ionomycin. **g**, Proliferation of GPR183^A+^ cells and control NK-92 cells quantified via decay of cell-trace dye. Left, histogram plot. Right, relative mean fluorescence intensity (MFI) quantification (n = 3). The graphics in (**a**) were created with Biorender.com.

NK cells with GPR183 CRISPR activation (denoted as GPR183^A+^) overexpressed genes involved in NK cell cytotoxicity (e.g., *FASLG* encoding Fas ligand, *GZMB* encoding granzyme B, *PRF1* encoding for perforin, *ICAM1*, *IFNG*, *NFAT5*, *SH2D1B*, and *SH3BP2*) and chemokines (*CCL18*, *CCL3*, *CCL3L1*, *CCL4*, *CCL4L2*, *CCL5;* **Figure 3b,c**), suggesting that *GPR183* CRISPR activation may also support NK cell cytotoxicity and ability to recruit other immune cells to the tumor, respectively. Aligned with GPR183 ligand being a form of oxidized cholesterol (7α,25- OHC), the genes downregulated in GPR183^A+^ cells were enriched for cholesterol and sterol biosynthesis genes (*P* < 1×10^-10^, e.g., *ACLY, CNBP, CYP51A1, DHCR7, FDFT1, FDPS, HMGCS1, IDI1, INSIG1, LSS, MSMO1, MVD, SQLE, SERBP1,* and *TM7SF2*), as well as oxidative phosphorylation, chromatin organization and other gene sets (**Figure 3b,c** and **Supplementary Table 4**).

These transcriptional readouts indicated that GPR183 does not disrupt and may enhance NK cell cytotoxicity and effector functions, as we turned to examine via functional readouts. Aligned with the transcriptional data, GPR183^A+^ NK cells show a more specific and efficacious response to stimuli compared to control NK cells. GPR183^A+^ NK cells exhibited increased IFNγ secretion and cytotoxicity in coculture with MDA-MB-231 breast cancer cells (**Figure 3d**). Upon PMA/ionomycin stimulation, GPR183^A+^ NK cells also show a significant increase in degranulation, as indicated by the surface expression of CD107a – a membrane-bound molecule commonly used as a proxy for cytotoxic degranulation – and IFNγ production (**Figure 3e,f**). Interestingly, MDA-MB-231 cells express the genes encoding oxysterol and 7α,25-OHC synthesizing enzymes and overexpress them in response to IFNγ and TNF (**Supplementary Figure 6c**), indicating a potential positive regulation across GPR183^A+^ NK and cancer cells. Importantly, GPR183^A+^ cells show minimal degranulation and IFNγ production at baseline (**Figure 3e,f**) and proliferate at similar rates to those of control NK cells, as quantified via cell-trace dye assays over 4 days (**Figure 3g**). These features are particularly important due to the potential risks associated with cell therapies, including cytokine release syndrome (48) and malignant transformation of the transferred immune cells (49).

Given these findings, we focused on GPR183 to examine whether metabolite-sensing GPCRs can make engineered lymphocytes spatially targeted and more efficacious in vivo.

### GPR183 activation in NK and CD8 T cells enhances directional migration to breast cancer

Given the biochemical and biophysical properties of GPR183, its association with tumor infiltration in multiple cell types, including CD8 T cells (**Figure 1e**, **Supplementary Figure 3a,b**), and the high degree of transcriptional similarity between NK cells and CD8 T cells, we turned to investigate GPR183-mediated directional migration in NK-92, as well as human primary NK cells and human primary CD8 T cells. In vitro, GPR183^A+^ NK-92 showed a significant increase in cell migration to MDA-MB-231 supernatant and breast cancer tumor lysate compared to control NK- 92 cells (**Figure 4a,b**). To test GPR183 in primary NK cells, we isolated and expanded primary NK cells from human peripheral blood mononuclear cells (PBMCs) using bead-based isolation and a feeder-cell free expansion method (50), which resulted in highly enriched CD56^+^CD3^-^ NK cells at > 92% (**Supplementary Figure 7a**). We then transduced the primary NK cells using spin-infection with high titer with either GPR183 ORF or control lentivirus (**Figure 4c**). As observed in NK-92 cells, GPR183^A+^ primary NK cells exhibited significantly increased migration to 7α,25- OHC compared to control NK cells from the same donor, which were transduced with the control lentivirus (**Figure 4d**). To test GPR183 in primary CD8 T cells, we isolated CD8 T cells from human PBMCs (**Supplementary Figure 7b**) and transduced them with either GPR183 ORF or control lentivirus (**Figure 4e**). GPR183^A+^ primary T cells displayed increased migration to 7α,25- OHC (**Figure 4f**), MDA-MB-231 supernatant (**Figure 4g**), and breast cancer tumor lysate (**Figure 4h**) compared to control CD8 T cells from the same donor, which were transduced with the control lentivirus. Focusing on NK-92 for in vivo testing, we find that GPR183^A+^ NK-92 cells intravenously transferred into MDA-MB-231 xenograft-bearing mice are significantly more abundant in the tumors compared to control NK-92 cells (**Figure 4i** and **Supplementary Figure 8**).

**Figure 4.**
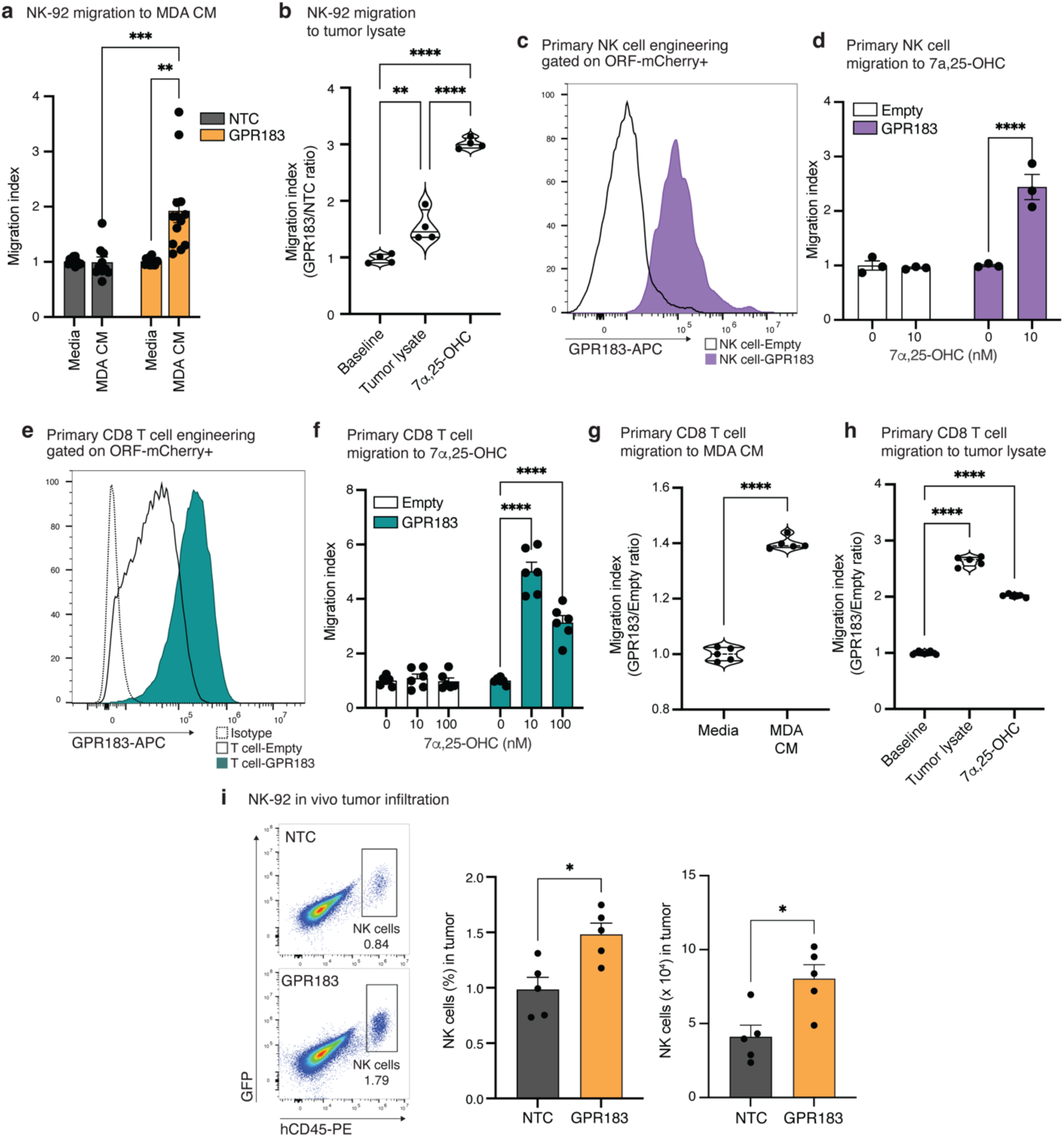
GPR183 activation enhances NK and T cell migration to breast cancer. **a**, GPR183^A+^ NK-92 cells show a significant increase in migration to MDA-MB-231 cell conditioned media compared to control NK-92 cells based on transwell migration assay with flow cytometry cell count readouts (n = 10 for NTC and n = 13 for GPR183^A+^). MDA CM: MDA-MB-231 conditioned media. **b**, GPR183^A+^ NK-92 cells show a significant increase in migration to tumor lysates compared to control NK-92 cells in competitive transwell migration assays with flow cytometry cell count readouts (n = 4 per group; **Methods**). **c**, GPR183 cell surface expression in GPR183^A+^ and control primary human NK cells. **d**, GPR183^A+^ primary NK cells show a significant increase in migration to 7α,25-OHC compared to control primary NK cells in a transwell migration assay with flow cytometry cell count readouts (n = 3 per group). **e**, GPR183 cell surface expression in GPR183^A+^ and control primary human CD8 T cells. (**f-h**) GPR183^A+^ primary T cells show a significant increase in migration to 7α,25-OHC (**f**), MDA-MB-231 cell conditioned media (**g**), and tumor lysates (**h**) in transwell migration (**f**) and competitive transwell migration assays (**g-h**) with flow cytometry cell count readouts [n = 6 per group in (**f**) and n = 5 per group in (**g**) and (**h**)]. **i**, GPR183^A+^ NK-92 cells show a significant increase in infiltration to breast cancer tumors in vivo (n = 5 per group), as shown based on representative flow cytometry plots (left), percentage (middle), and number of NK cells in the tumor (right). 100 nM 7α,25-OHC was used as a positive control in competitive migration assay in (**b**) and (**h**). Three independent experiments were pooled for (a). Data are representative of two independent experiments for (**b** and **f**-**h**) and are presented as the mean ± s.e.m. for (**a**, **d**, **f**, and **i**). Data analysis was performed using one-way ANOVA with Dunnett’s multiple comparisons for (**a**, **b**, **f**, and **h**) and two-tailed unpaired Students’ t-test for (**d**, **g**, and **i**) (\*\*\*\**P* < 0.0001 and \*\*\**P* < 0.001).

### GPR183 activation in NK-92 and CAR NK-92 results in better breast cancer tumor control

As insufficient recruitment and infiltration are a limiting factor of cell therapies in solid tumors, we turned to examine the efficacy of GPR183^A+^ NK cells in controlling breast cancer tumors in vivo (**Figure 5a**). First, intravenous transfer of GPR183^A+^ NK-92 cells to NSG female mice bearing MDA-MB-231 xenografts significantly delayed tumor growth compared to both untreated mice (i.e., treated with cell-free media) and mice treated with control NK cells (i.e., NK-92 cells transduced with control lentivirus; **Figure 5b**), such that the same regimen of control NK-92 cell transfer had no impact on tumor growth. Second, to test whether GPR183 activation can enhance the in vivo efficacy of CAR NK cells, we cloned a CAR construct that targets EpCAM (epithelial cell adhesion molecule) on the surface of breast cancer cells (**Figure 5c** and **Supplementary Figure 9a**) and co-introduced the construct (anti-human EpCAM-CAR-GFP) with GPR183-ORF- mCherry or empty-ORF-mCherry into NK-92 cells (**Supplementary Figure 9b**), resulting in antigen-specific cell cytotoxicity (**Supplementary Figure 9c,d**). Intravenous transfer of GPR183^A+^ CAR-NK-92 showed significantly better control of MDA-MB-231 tumor growth compared to both the untreated group (i.e., treated with cell-free media) and the group treated with control (i.e., empty-ORF-mCherry) CAR-NK-92 cells, demonstrating the synergistic effect of CAR and GPR183 (**Figure 5d** and **Supplementary Figure 9e**). Together, these experiments demonstrate that engineering NK cells to express GPR183 provides a mechanism to enhance NK cell antitumor efficacy in vivo.

**Figure 5.**
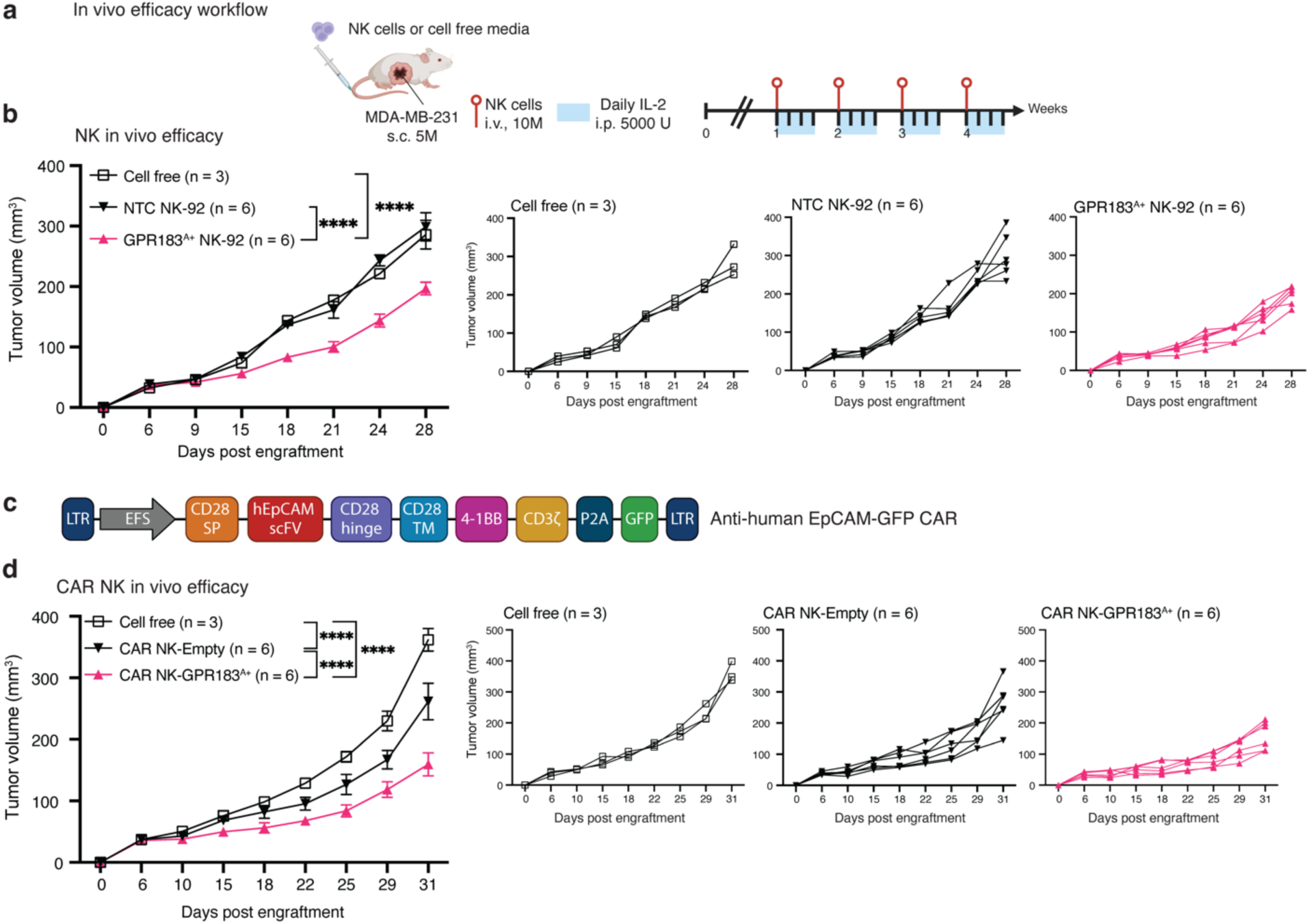
Engineering NK-92 and CAR NK-92 to express GPR183 significantly improves NK-mediated tumor control in vivo. **a**, Disease outcome study design for (**b**) and (**d**). **b**, Growth curve of MDA-MD-231 breast cancer tumors in NSG mice treated with cell-free media (n = 3 mice), control NK-92 (n = 6 mice) or GPR183^A+^ (n = 6 mice) NK-92 cells. **c**, EpCAM CAR construct. **d**, Growth curve of MDA-MD-231 breast cancer tumors in NSG mice treated with cell-free media (n = 3 mice), anti-EpCAM-CAR-NK-92 (n = 6 mice), or GPR183^A+^ anti-EpCAM- CAR-NK-92 (n = 6 mice). The results from a biological replicate of this experiment (n = 13) are depicted in **Supplementary Figure 9e**. In (**b**) and (**d**) data are shown as the mean ± s.e.m. (left) and individual growth curves per mouse (right); statistical significance was evaluated using a two-way ANOVA with Sidak correction for multiple hypotheses testing (\*\*\*\**P* < 0.0001, \*\*\**P* < 0.001, \*\**P* < 0.01 and \**P* < 0.05). The graphics in (**a**) and (**c**) were created with Biorender.com.

## DISCUSSION

Identifying mechanisms to redirect immune cells to specific sites in the body and within a target tissue is critical. Here, we identified an underappreciated and programmable mechanism to recruit cytotoxic lymphocytes to breast cancer tumors via activation of metabolite-sensing GPCRs. Focusing on GPR183, we show that it redirects NK and T cells to 7α,25-OHC, breast cancer cells, and breast cancer tumors and significantly improves NK-mediated tumor control. Our study thus demonstrates that metabolic sensing can be rewired and used to recruit cytotoxic lymphocytes to solid tumors, opening a new avenue for the development of spatially targeted cell therapies.

Altered metabolism has been widely recognized as a hallmark of cancer (21), allowing cancer cells to generate biomass and sustain anabolic processes critical for tumor proliferation. Alongside the Warburg effect (which was first described in 1931 (19,20)), altered lipid metabolism is among the most prominent metabolic alterations in cancer (22). These unique metabolic properties of tumors have provided means for cancer detection via Positron Emission Tomography (PET) imaging, using radiolabeled glucose analogs like fluorodeoxyglucose and, more recently, also PET tracers targeting lipid metabolism (as choline (51) and acetate (52) analogs). Such metabolic reprogramming not only supports tumor growth and allows cancer detection for diagnostics but has also been reported to exert immune regulatory effects. For instance, aberrant lipid metabolites enriched in tumors, such as prostaglandins, lysophosphatidic acid, and cholesterols, have been shown to suppress the antitumor effects of cytotoxic CD8 T cells and NK cells (45,53–58). In contrast, here we demonstrate that these metabolic attributes can be exploited to mobilize cytotoxic lymphocytes to the tumor via exogenous expression of metabolite-sensing GPCRs, resulting in better tumor control.

This, of course, raises the concern that activation of such metabolite-sensing GPCRs, while beneficial for recruitment, may suppress cytotoxic effector functions, as recently shown for GPR34 (45). Based on our data, GPR183 is not suppressing NK cell effector functions and may further enhance such functions upon stimuli. While additional investigation is needed, we note that GPR183, as well as all other seven hits identified in our screen, are coupled with Ga_i/o_ protein signaling that suppresses cAMP production (59,60), thus counteracting the GPCR Ga_s_-cAMP pathway that has been recently shown to promote CD8 T cell dysfunction (59).

We also note that while chemokine receptor inhibition is likely to disrupt NK tumor homing, (over)expression of most chemokine receptors did not have a major impact on NK cell homing to breast cancer tumors in our screens, potentially because these genes are already expressed at sufficiently high levels by NK cells at baseline. Supporting this, the only chemokine receptor that improved NK tumor homing in our screens was *CXCR2*, a chemokine receptor that is rarely expressed by lymphocytes and is expressed mainly by neutrophils (61). This demonstrates the value and orthogonal view of gene function revealed by gain-of-function studies.

GPR183 is an oxysterol-responsive receptor broadly expressed across immune cells, including naïve B cells (42,62,63), plasma cells (64), naïve CD4 (65), innate lymphoid cells (66), and dendritic cells (67). GPR183 orchestrates the positioning of the cells within lymphoid and mucosal tissues, where its ligands are produced via oxysterol-synthesizing enzymes. Inflammation-induced upregulation of these enzymes amplifies ligand availability, thereby enhancing the recruitment of GPR183-expressing cells to inflamed sites (66,68,69). Interestingly, a recent study demonstrated that GPR183 knockout impairs homing of naïve B and a subset of memory T cells into inflamed lymph nodes and inflamed nonlymphoid endothelium of tumors in an oxysterol-dependent manner (68). Here, we demonstrate that GPR183 overexpression in NK and CD8 T cells enhances their trafficking to breast cancer tumors and factors secreted by the breast cancer cells themselves. We show that engineering NK and CAR NK cells to express GPR183 augments in vivo efficacy, demonstrating this recruitment mechanism can form a potential basis for spatially targeted cell therapies.

Further investigation will be needed to fully elucidate the cancer-immune circuits, ligands, and therapeutic potential of *GPR183*, *GPR84*, *GPR34*, *GPR18*, *FPR3*, and *LPAR2*, as well as the potentially synergistic effect of their combined activation. As shown here, these receptors provide means to mobilize lymphocytes to breast cancer tumors in ways that diverge and may synergize with or reinforce chemokine-based chemoattraction. Taken together, we anticipate that this study will provide a new lens through which to investigate and leverage cancer abnormal metabolism and open a new line of investigation to activate metabolite sensing GPCRs to improve tumor-targeted cell delivery and mobilize tumor-reactive cytotoxic lymphocytes to solid tumors.

## ACKNOWLEDGMENTS

We thank M. Tsai for his help with setting up models related to this work and Dr. Michael C. Bassik for providing the HTLA cell line. Y.K. is a recipient of the Stanford School of Medicine Dean’s Postdoctoral Fellowship and of the Basic Science Research Program fellowship (RS-2023- 00244834) through the National Research Foundation of Korea (NRF), funded by the Ministry of Education. L.J. is a Chan Zuckerberg Biohub Investigator and an Allen Distinguished Investigator. L.J. holds a Career Award at the Scientific Interface from the Burroughs Wellcome Fund (BWF) and a Liz Tilberis Early Career Award from the Ovarian Cancer Research Alliance (OCRA). This study was supported by BWF (1019508.01; L.J.), National Human Genome Research Institute (NHGRI; U01HG012069; L.J.), Under One Umbrella, Stanford Women’s Cancer Center, Stanford Cancer Institute, a National Cancer Institute (NCI)-designated Comprehensive Cancer Center (311982; L.J), as well as funds from the Departments of Genetics at Stanford University (L.J.).

## AUTHOR CONTRIBUTIONS

Y.K. and L.J. designed the study. Y.K., R.V.A., C.S., and O.L. performed the experiments under L.J. supervision. Y.K. and L.J. visualized and analyzed data. Y.K. and L.J. wrote the paper. L.J. obtained funding. L.J. supervised the study. All authors reviewed and approved the paper.

## ADDITIONAL INFORMATION

### Extended data is available

**Supplementary Tables** are provided as separate Excel files.

**Supplementary Table 1.** sgRNA library information, including target gene annotation and differential expression in breast cancer tumors vs. blood samples (a) and spacer sequence per sgRNA (b).

**Supplementary Table 2.** In vivo CRISPR activation screen MAGeCK summary statistics (a) and output per screen (b-g).

**Supplementary Table 3.** In vitro CRISPR activation screen MAGeCK summary statistics (a) and outputs per screen (b-h).

**Supplementary Table 4.** Perturb-seq DEGs (a-h) and GSEA (i-k).

**Supplementary Table 5.** Gene sequences for ORF, qPCR primer, and sgRNA.

---

**Supplementary Figure 1.**
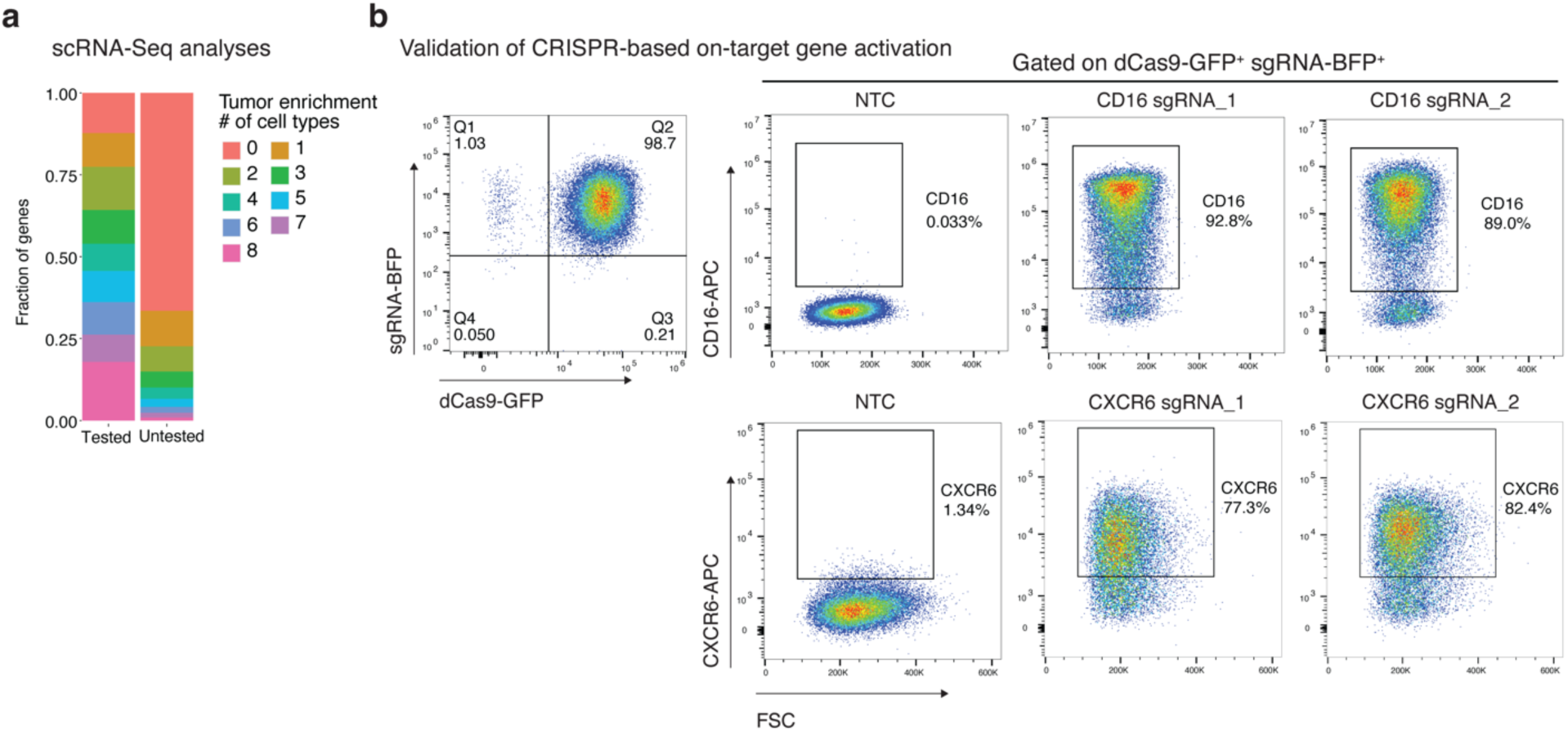
Experimental design of CRISPR activation screens in NK-92 cells. **a**, CRISPR activation sgRNA library design based on scRNA-Seq data: Genes stratified by the number of cell subtypes in which they were found to be significantly overexpressed in the tumor compared to blood samples of breast cancer patients (27), shown for the genes tested (left) or untested (right) in the screens. **b**, Validations of CRISPR/dCas9-SAM system in NK-92 via flow cytometry of target gene protein expression of the cell surface for two different sgRNAs targeting *FCGR3A* (encoding for CD16; top) and *CXCR6* (bottom).

**Supplementary Figure 2.**
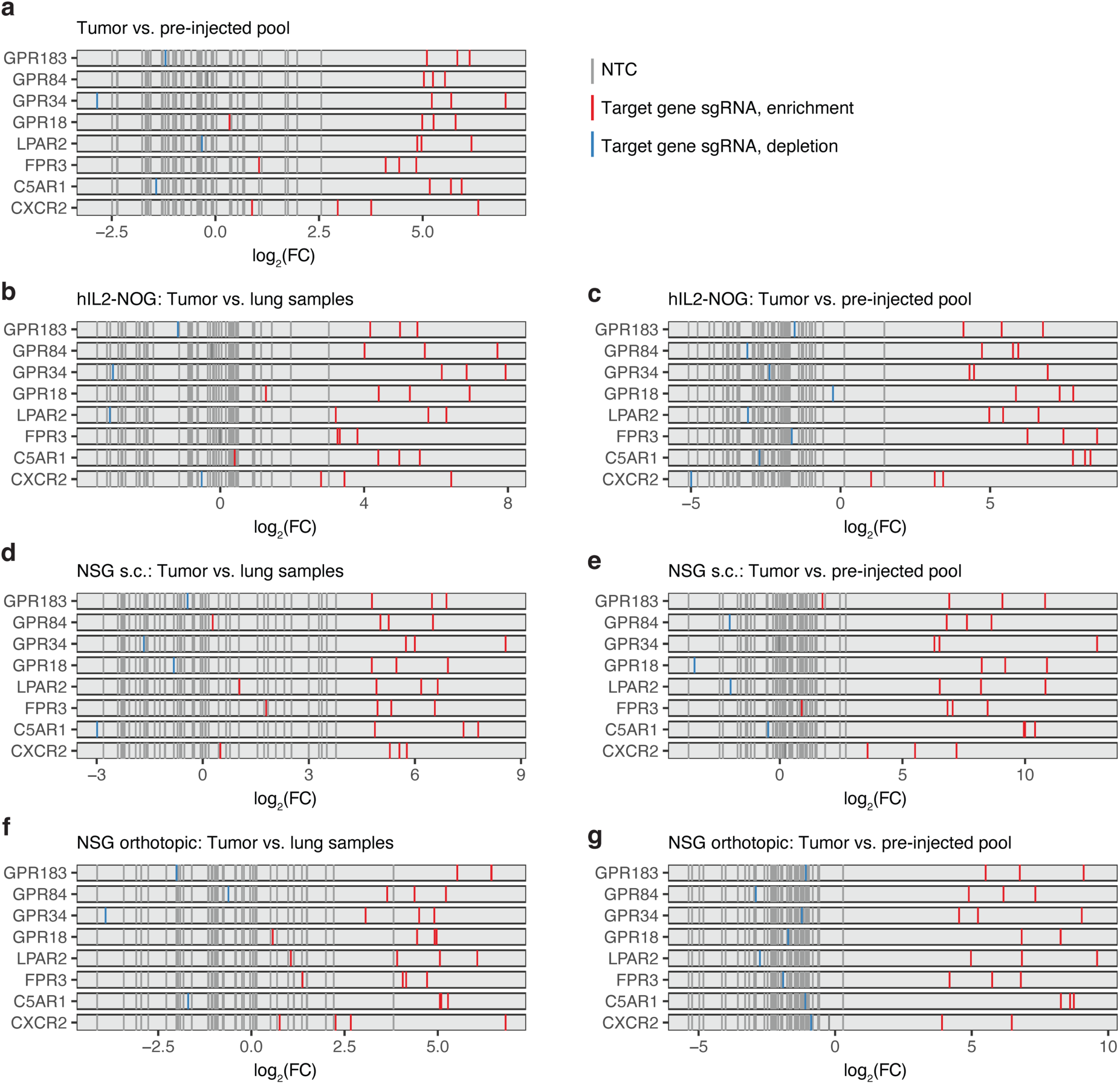
Tumor infiltration hits identified in vivo across different models. (**a-g**) Fold change of sgRNAs in tumor vs. lung or pre-injected NK cell samples shown for sgRNAs targeting indicated genes. Combined analysis of all in vivo models when comparing tumor to pre-injected samples (**a**). In vivo screen in hIL-2-NOG mice (**b,c**) and NSG mice with subcutaneous MDA-MB-231 implantation (**d,e**), and in vivo screen in NSG mouse with orthotopic MDA-MB- 231 implantation (**f,g**).

**Supplementary Figure 3.**
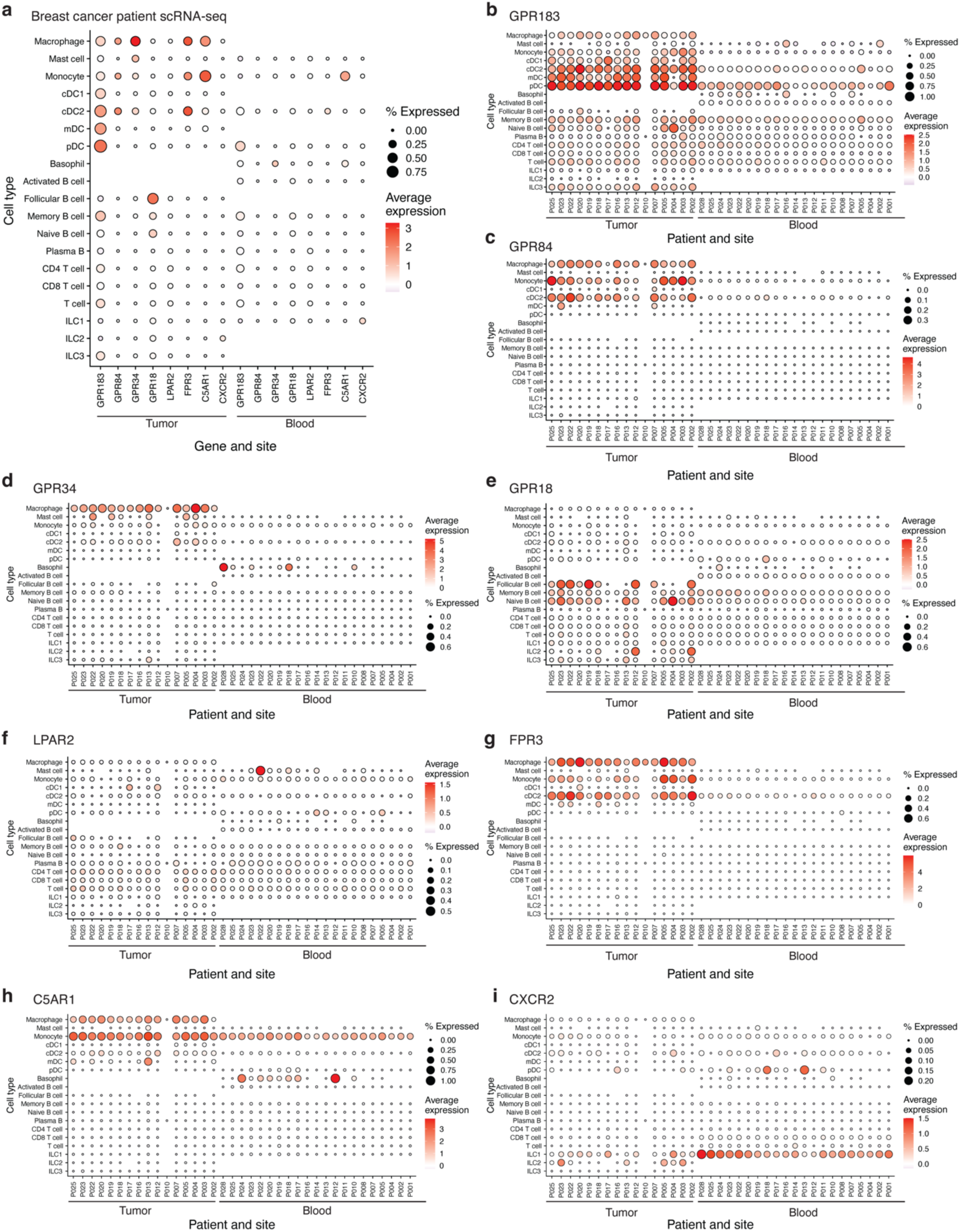
Hit expression in breast cancer patient scRNA-seq data. **a**, Expression of hits in the tumor and blood samples per cell subtypes. (**b-i**) *GPR183* (**b**), *GPR84* (**c**), *GPR34* (**d**), *GPR18* (**e**), *LPAR2* (**f**), *FPR3* (**g**), *C5AR1* (**h**), and *CXCR2* (**i**) expression, stratified by cell type, patient and site (blood or tumor). cDC1: conventional type 1 dendritic cells; cDC2: conventional type 2 dendritic cells; mDC: mature DC; pDC: plasmacytoid DC.

**Supplementary Figure 4.**
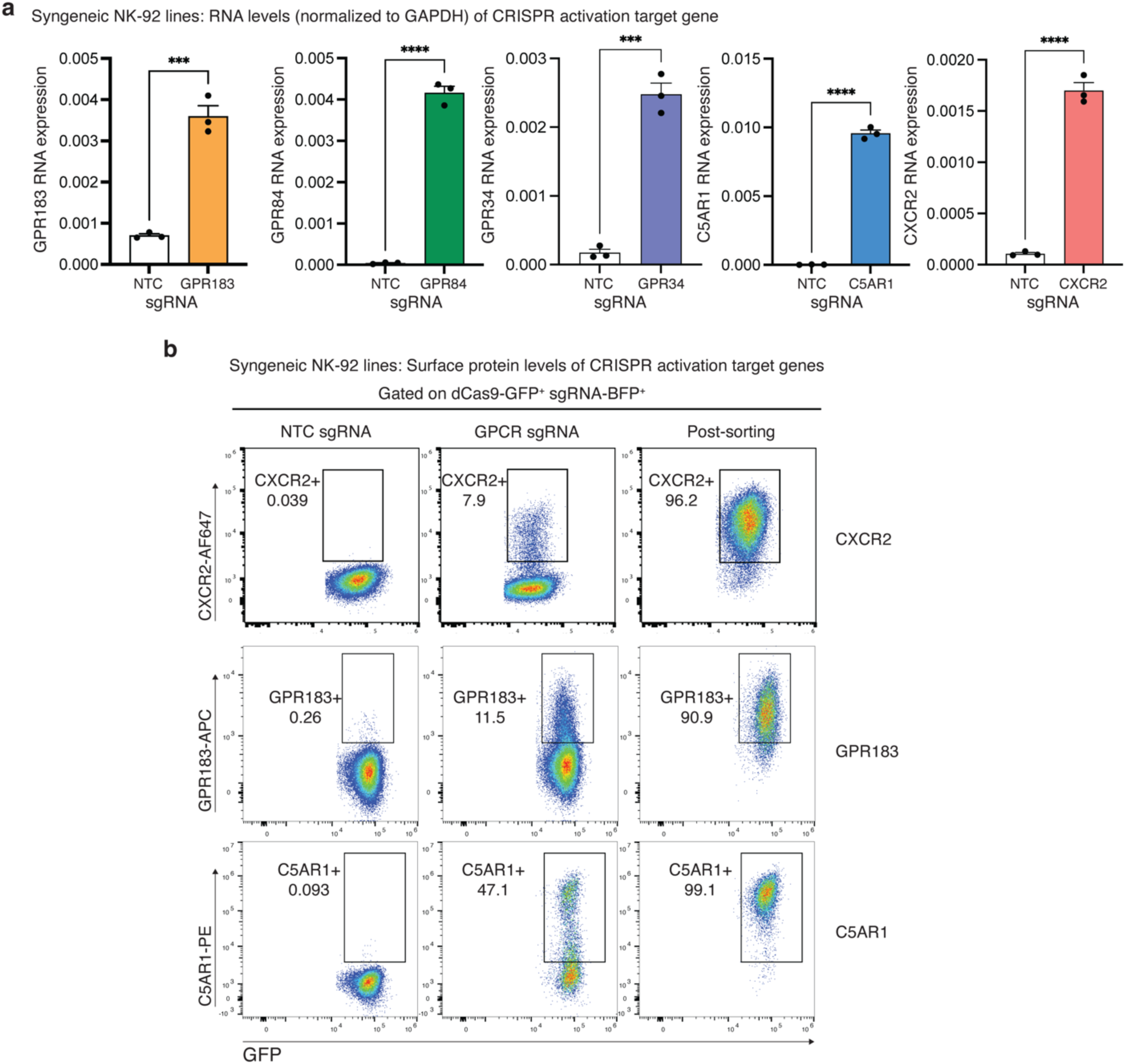
Syngeneic NK-92 cell lines with constitutive expression of top hits. **a**, qPCR quantification of *GPR183*, *GPR84*, *GPR34*, *CXCR2*, and *C5AR1* expression in control NK-92 cells and upon CRISPR activation of the target GPCRs (n = 3 per group). **b**, Surface expression levels of CXCR2, GPR183, and C5AR1 in NK-92 cells post-transduction with a NTC sgRNA (left) or with the target gene sgRNAs (middle) and 5 days after fluorescence-activated cell sorting (FACS) for surface expression of the target GPCR (right). Data are representative of two independent experiments and presented as the mean ± s.e.m. \*\*\*\**P* < 0.0001 and \*\*\**P* < 0.001, two-tailed unpaired Students’ t-test.

**Supplementary Figure 5.**
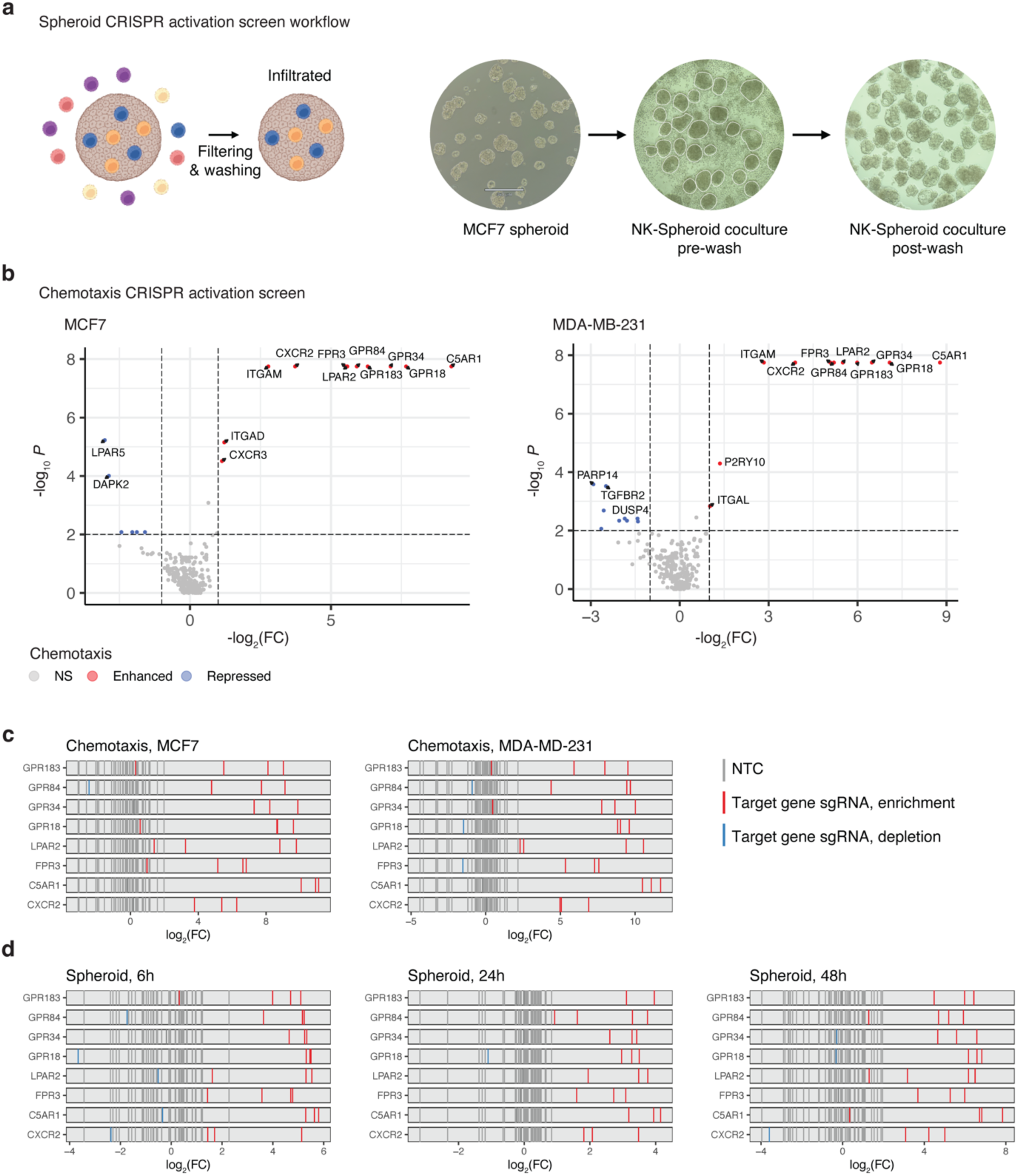
In vitro CRISPR activation screens. **a**, Experimental scheme and image of spheroid-NK coculture system. **b**, Chemotaxis NK-92 CRISPR activation screens: Significance (y-axis) and fold change (x-axis) of each target gene in enhancing (positive) or depleting (negative) migration to MCF7 (left) and MDA-MB-231 (right) supernatant. (**c,d**) Log transformed fold change of sgRNAs enriched in NK-92 cells chemoattracted to MCF7 (left) and MDA-MB-231 (right) supernatant and in MCF7 spheroids across different time points (**d**). The graphics in (**a**) were created with Biorender.com.

**Supplementary Figure 6.**
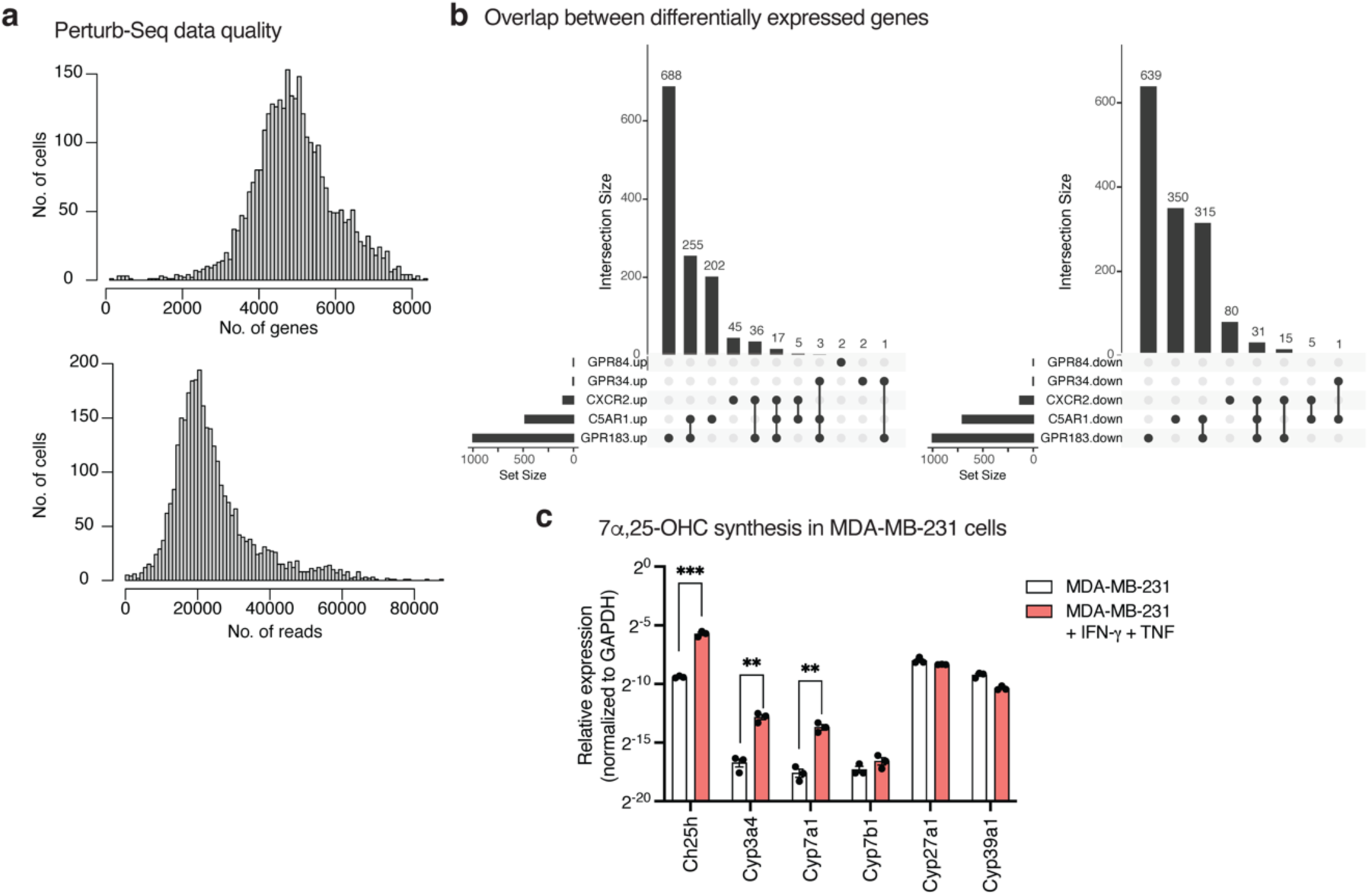
Perturb-seq and gene expression regulation. **a**, Number of genes (top) and reads (bottom) detected per cell in the Perturb-seq data. **b**, Overlap between the differentially expressed genes identified for perturbations that alter the NK-92 cell transcriptome in the Perturb-seq screen, shown separately for up (left) and down (right) regulation. **c**, Expression levels of 7α,25-OHC synthesizing enzymes in MDA-MB-231 cells in monoculture with and without IFNγ and TNF treatment (n = 3 per group). Expression levels were measured via qPCR and normalized to *GAPDH*. Data are representative of two independent experiments and presented as the mean ± s.e.m. \*\*\**P* < 0.001 and \*\**P* < 0.01, two-tailed unpaired Students’ t-test.

**Supplementary Figure 7.**
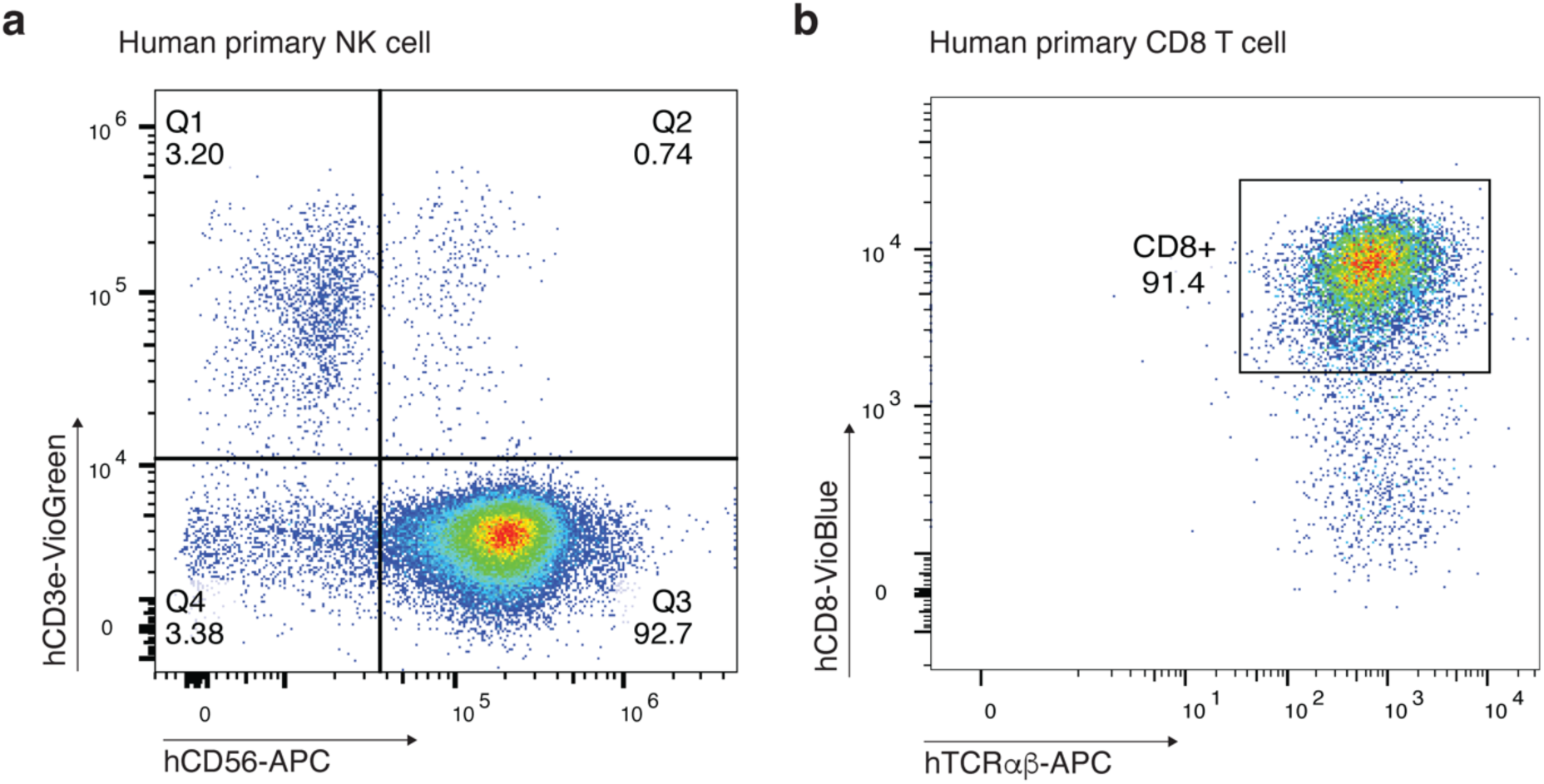
Human primary NK cells and CD8 T cell isolation. **a**, Purity of human primary NK cells based on the NK cell marker CD56. **b**, Purity of human primary CD8 T cells based on CD8a and human TCRαβ.

**Supplementary Figure 8.**
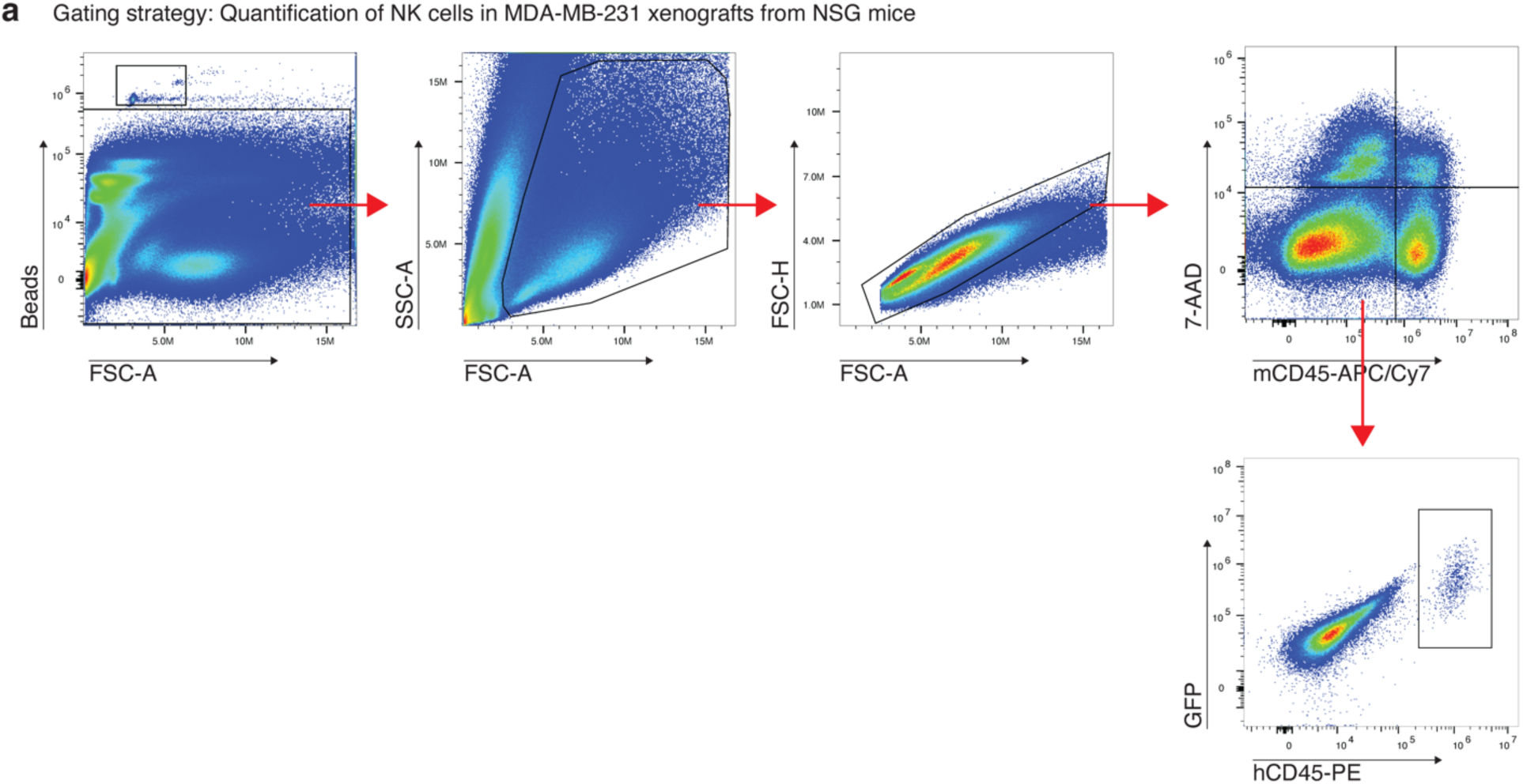
A gating strategy for quantifying tumor infiltrating NK-92 cells. **a**, Flow cytometry plots show the gating strategy to detect and quantify the number and relative abundance of NK-92 cells in the tumor. Only live (7-AAD negative) cells were considered. mCD45^-^**/**hCD45^+^/GFP^+^ were identified as NK-92 cells in the NSG mice. mCD45: mouse CD45; hCD45: human CD45; GFP: Green fluorescent protein.

**Supplementary Figure 9.**
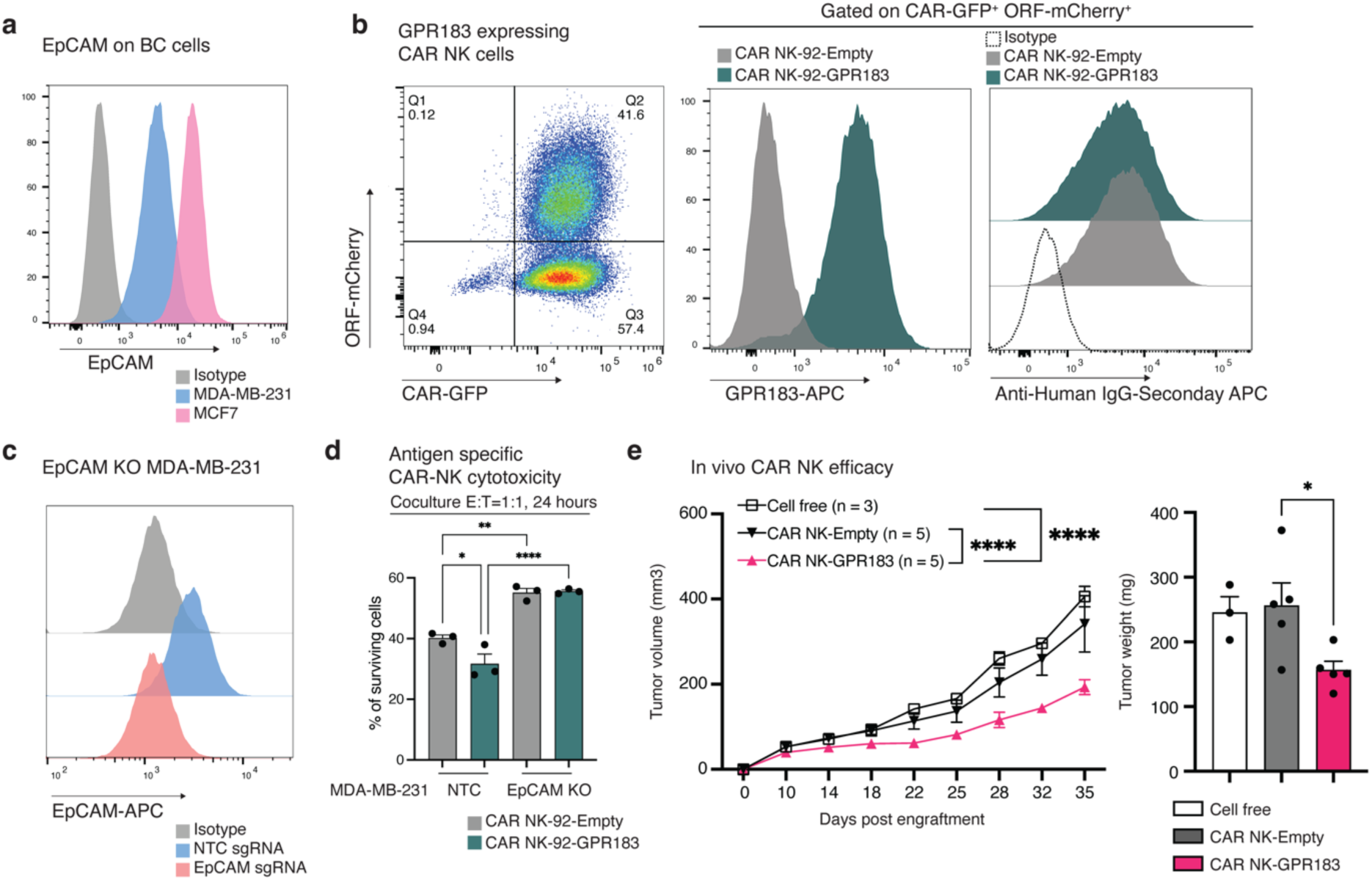
Generating and validating CAR NK-92 cells. **a**, EpCAM protein surface expression in MDA-MD-231 and MCF7 cells quantified via flow cytometry. **b**, FACS of anti-EpCAM-CAR^+^ and GPR183^A+^ NK-92 cells via mCherry and GFP, respectively (left), followed by confirmation of GPR183 surface expression via an anti-GPR183 antibody (middle) and confirmation of CAR expression via anti-human IgG (right). **c**, Flow cytometry measurements of EpCAM surface protein levels in control (NTC) and EpCAM KO MDA-MB-231 cells. **d**, Antigen-dependent function of control and GPR183^A+^ anti-EpCAM-CAR^+^ NK-92 cells, shown based on cytotoxicity in coculture with control (NTC, EpCAM wildtype) and EpCAM KO MDA-MB-231 cells at 1:1 E:T ratio for 24 hours (n = 3 per group). **e**, Another in vivo disease outcome experiment was conducted to examine the impact of GPR183 on CAR NK-92 cell efficacy in vivo and validate the results shown in Figure 5d. Left: Growth curve plots depict MDA-MD-231 tumors in NSG mice treated with cell-free media (n = 3 mice), anti-EpCAM-CAR-NK-92 (n = 5 mice), or GPR183^A+^-anti-EpCAM-CAR-NK-92 (n = 5 mice). Data are presented as the mean ± s.e.m. two-way ANOVA with Sidak correction for multiple hypotheses testing. Right: Tumor weight after tissue dissection at day 35 shown as the mean ± s.e.m. One-way ANOVA was performed with Dunnett’s multiple comparisons. \*\*\*\**P* < 0.0001, \*\*\**P* < 0.001, \*\**P* < 0.01 and \**P* < 0.05.

## METHODS

### Study approval

The study received institutional regulatory approval. All experiments involving recombinant DNA and other biosafety considerations were performed under the guidelines and with the approval of Stanford University Environment, Health and Safety Committee (APB-3910-L.J.0921; APB- 5416-L.J.0724). All experiments involving materials from human samples were performed under the guidelines and with the approval of Stanford University Institutional Review Board (protocols IRB-13942). All animal experiments were performed under guidelines and with the approval of the Stanford University Institutional Animal Care and Use Committee (IACUC; APLAC-34218).

### Cell lines

NK-92 (CRL-2407), MDA-MB-231 (HTB-26), and MCF7 (HTB-22) cells were purchased from ATCC and Lenti-X 293T cells (632180) were bought from Takara Bio. NK-92 cells were cultured with RPMI 1640 medium GlutaMAX™ supplement (Gibco, 61870036) containing 5 mM MEM non-essential amino acids solution (cytiva, SH30598.01), 5 mM Sodium pyruvate 100mM solution (Cytiva, SH30239.01), 10% fetal bovine serum (Gibco, A3840102), 1% penicillin/streptomycin (referred here as NK base media), and 200 U/ml IL-2 (Frederick National Laboratory). MDA-MB- 231, MCF7, and Lenti-X 293T cells were cultured with DMEM medium containing 5 mM MEM non-essential amino acids solution, 5 mM Sodium pyruvate 100mM solution, 10% fetal bovine serum (FBS), and 1% penicillin/streptomycin. All cell lines were maintained at 37 °C in 5% CO_2_ and tested for mycoplasma by PCR. All cell lines were authenticated through ATCC using short tandem repeat profiling.

### Lentivirus production

Lentivirus was generated from low-passage Lenti-X 293T cells as follows. Lenti-X 293T cells were seeded in Opti-MEM I reduced serum medium (Gibco, 31985088) containing 1X GlutaMAX supplement, 1X sodium pyruvate, 1X MEM non-essential amino acids, and 5% FBS (cOpti-MEM) in a T75 flask at 80-90% confluency. The following day, 14 μg transfer vector, 10 μg psPAX2 (Addgene, 12260), 4.33 μg pMD2.G (Addgene, 12259), and 85 μl MirusBio TransIT-Lenti (MirusBio 6603) were mixed in 2.8 ml of pre-warmed Opti-MEM. After incubation at room temperature for 10 minutes, the mixture was added to the cells. After 6 hours of transfection, the medium was replaced with fresh cOpti-MEM plus 1X ViralBoost (Alstem Bio cat, VB100). Viral supernatant was collected at 48 hours post-transfection, filtered using 0.45-μm filters, and concentrated using Lenti-X Concentrator (Takara Bio, 631232). The virus was aliquoted and stored at −80 °C.

### Custom CRISPR activation library of NK-92 cells

Protospacer sequences of the custom activation library were obtained from the Human Genome-Wide CRISPRa-v2 Libraries and Human CRISPR Activation Library (29). The pooled sgRNA library was cloned into pKVL2-U6gRNA_SAM (BbsI)-PGKpuroBFP-W vector (Addgene, 112925) by Genscript. In total, the library consists of 1,070 sgRNAs targeting 256 genes with four guides per gene and 46 NTCs (**Supplementary Table 1**). Lentivirus was prepared as described above. For applying the CRISPR SAM system in NK-92 cells, NK-92 were sequentially transduced with lentivirus encoding dCas9-VP64-GFP (Addgene, 61422) and lentiMPH V2 (Addgene, 89308) at a multiplicity of infection (MOI) < 0.3. Transduced NK-92 cells were selected by GFP-based cell sorting and 500 μg/ml hygromycin B Gold (InvivoGen, ant-hg-1). The dCas9-MPH NK-92 cells were transduced with the lentiviral custom sgRNA library at a MOI < 0.3 and selected with 1 μg/ml puromycin for 7 days.

### *In vivo* NK CRISPR activation screen

Three types of in vivo screens were conducted using the following models: (1) NOD.Cg-Prkdcscid Il2rgtm1Sug Tg (CMV-IL2)4-2Jic/JicTac (Taconic Biosciences, hIL-2-NOG, 13440-F) mice with subcutaneous breast cancer xenografts, (2) NOD.Cg-Prkdcscid Il2rgtm1Wjl/SzJ (The Jackson laboratory, NSG, 005557) mice with subcutaneous breast cancer xenografts, (3) NSG mice with orthotropic mammary fat pad breast cancer xenografts. Seven to eight-week-old female NSG and hIL-2-NOG mice were purchased from Jackson Laboratory and Taconic Biosciences, respectively. All xenografts resulting from 5 x 10^6^ MDA-MB-231 cells that were injected (subcutaneously into the right flank or orthotopically into the left inguinal mammary fat pad) in a 1:1 mix of serum-free DMEM and Matrigel (Corning, 354483). After two weeks, when tumor size reached ∼100 mm^3^, 10^7^ library NK cells (i.e., dCas9-MPH NK-92 transduced with the sgRNA library) were adoptively transferred into tumor-bearing mice by intravenous tail-vein injections. For the experiment with NSG mice, human IL-2 (5000 U) was injected intraperitoneally immediately before NK cell transfer. Twenty-four hours after NK infusion, mice were euthanized, and tumors and lungs were collected and immediately processed for genomic DNA (gDNA) extraction.

### Chemoattraction NK CRISPR activation screen

Library NK cells (i.e., dCas9-MPH NK-92 transduced with the sgRNA library) were harvested and washed with serum-free RPMI medium. Cells were resuspended in RPMI containing 1% FBS and counted using Countess 3 Automated Cell Counter (Invitrogen, AMQAX2000). 2.5 x 10^6^ cells were added on a 24-mm transwell with 8 µm pore (Corning, 3428) in 2ml RPMI containing 1% FBS. Next, 2.7 ml MDA-MB-231 and MCF7 cancer cell conditioned media was added to the bottom plate and incubated at 37 °C and 5% CO_2_ for 4 hours. After incubation, the NK cells that reached the bottom plate were collected and immediately processed for gDNA extraction. Cells from 6 wells were pooled for a single experimental replicate, and three replicates were collected per condition.

### Spheroid NK CRISPR activation screen

For a scalable spheroid generation, MCF7 cells were seeded at 1 x 10^6^ cells/ml in 10 ml of DMEM containing 1% methylcellulose (Sigma-Aldrich, H7509) in a 90 mm ultra-low attachment dish (Sbio, MS-90900Z) and incubated for 2 days for spheroid formation. MCF7 spheroids were collected, diluted with PBS, centrifuged, and resuspended in NK base media. A small fraction of cells was taken, digested with TrypLE (Gibco, 12605010), and counted. 5 x 10^6^ custom library NK cells and 2.5 x 10^6^ MCF spheroids were seeded at 2:1 E:T in 12.5 ml of RPMI + IL-2 (200 U/ml) in 90 mm ultra-low attachment dish. The cocultures were harvested after 6, 24, and 48 hours and then filtered through a 70 μm strainer to separate non-infiltrated NK cells and spheroids. The filter was washed with PBS five times to remove residual non-infiltrated single cells. The remaining cells on top of the filter were collected and considered spheroid-infiltrating cells. Input NK cell pool was also in vitro cultured and collected at the same time points to obtain control reference samples. gDNA of spheroids and control samples were immediately processed. Cells from 6 dishes were pooled for a single experimental replicate, and three replicates were collected per condition.

### Genomic DNA extraction, sgRNA amplification, and sequencing

Genomic DNA of the cells obtained from in vitro culture, post-transwell migration, or spheroid screens was extracted using QIAamp DNA Mini or Midi Blood kit (Qiagen, 51104 or 51183), following manufacturer’s protocol. For in vivo samples, tumor and lung tissues were minced into small pieces, digested with Collagenase type IV (Gibco, 17104019) and DNase I (Roche, 11284932001) for 45 min at 37 °C with agitation, and then filtered through a 100 μm cell strainer to generate single-cell suspensions. Cells were pelleted and processed with a QIAamp DNA Midi Blood kit for gDNA extraction. The amount of gDNA was measured by Qubit and 5 μg of gDNA was used for the first-round PCR (5 μg per reaction; 2∼4 reactions per sample). Primers and thermocycler conditions for the first PCR: Forward: 5’-caaggctgttagagagataattagaattaatttgactgtaaacac-3’ and reverse: 5’-ccctacccggtagaattgg atccaaaaaaa-3’, 98 °C for 2 min, 18 cycles of (98 °C for 30 sec, 59 °C for 30 sec, 72 °C for 45 sec), and 72 °C for 3 min. The first PCR products were pooled and used as a template for the second-round PCR (5 μl per reaction; 2 reactions per sample) to add Illumina sequencing barcodes. For PCR2, thermocycler conditions were: 98 °C for 2 min, 11∼25 cycles of (98 °C for 10 sec, 63 °C for 10 sec, 72 °C for 25 sec), and 72 °C for 2 min. All PCR reactions were performed using NEBNext High-Fidelity 2X PCR master mix (NEW ENGLAND Biolabs, M0541L). The final products were gel purified, quantified with Qubit and BioAnalyzer, and then pooled for multiplexed sequencing. The pooled libraries were sequenced with 20 % PhiX control using Illumina MiSeq V3 or NextSeq 550 Mid Output sequencers.

### PRESTO-Tango assay

HTLA cells, a HEK293 cell line stably expressing a tTA-dependent luciferase reporter and a β- arrestin2-TEV fusion gene, previously developed by R. Axel (70) and used in the PRESTO-tango assay (43), were a gift from the laboratory of M. Bassik. HTLA cells were maintained in DMEM supplemented with 10% FBS, 1% penicillin/streptomycin, 2 μg/ml puromycin (InvivoGen, ant-pr-1), and 100 μg/ml hygromycin B at 37 °C and 5% CO_2_. For GPCR cloning, gene fragments encoding ORFs of GPCR hits (*CXCR2*, *C5AR1*, *GPR34*, *GPR84*, and *GPR183*; sequences provided in **Supplementary Table 5**) and TEV-V_2_ tail (encoding TEV-cleavage site and tTA transcription factor) were synthesized from Twist Bioscience and cloned into pLV-EF1a-IRES-Blast vector (Addgene, 85133), digested with BamHI-HF (NEW ENGLAND Biolabs, R3136) and EcoRI-HF (NEW ELGLAND Biolabs, R3101), with Gibson assembly. Cloned GPCR constructs were packaged with lentivirus and used to transduce HTLA cells. Transduced cells were selected with 8 μg/ml blasticidin (In vivoGen, ant-bl-05). For the PRESTO-tango assay, 4 x 10^4^ of each GPCR reporter and control HTLA cells were seeded in a white-walled, clear-bottom 96 well plate (Corning, 354651). One day after cell seeding, varying doses of relevant ligands, as indicated in the figures and figure legends, were added to each well and incubated at 37 °C and 5% CO_2_ for 24 hours. The next day, Bright-Glo solution (Promega, E2610) was added and incubated for 5 min at room temperature. Luminescence was measured using a Tecan infinite M1000 plate reader.

### Generating syngeneic NK-92 lines with GPCR overexpression

Syngeneic NK-92 lines with constitutive expression of top hits were generated as follows. Protospacer sequences of three NTC sgRNAs and three sgRNAs targeting top hits were obtained from Human Genome-Wide CRISPRa-v2 Libraries (71) and synthesized from IDT DNA. Oligo sequences are listed in **Supplementary Table 5**. The oligos were annealed and cloned into pKVL2-U6gRNA_SAM (BbsI)-PGKpuroBFP-W vector digested with FastDigest Bpil. Equal amounts of the three sgRNA vectors targeting a single GPCR were pooled and used for transfecting Lenti-X 293T cells to generate lentivirus. CRISPR/dCas9-MPH expressing NK-92 cells were transduced with sgRNA at MOI < 0.3 and selected with 1 μg/ml puromycin. CXCR2-, C5AR1-, and GPR183-engineered NK cells were further enriched with fluorescence-activated cell sorting based on protein expression. For ORF-based overexpression, gene fragments encoding GPR34 and GPR183 ORFs were synthesized from Twist Bioscience and cloned into pLV-EF1a-IRES-Puro vector (Addgene, 85132), digested with BamHI-HF and EcoRI-HF, with Gibson assembly. Cloned GPCR ORF constructs were packaged with lentivirus and used to transduce NK-92 cells. Transduced cells were selected with 1 μg/ml puromycin. In some experiments, the backbone was switched to pLV-mCherry (Addgene, 36084) and digested with BamHI-HF. The gene fragment sequences are listed in **Supplementary Table 5**. Engineered NK-92 cells were cryopreserved in liquid nitrogen until use. After thawing, the cells were cultured and passaged at least three times before being subjected to follow-up experiments, including in vitro and in vivo assays and Perturb-seq.

### In vitro transwell migration assays

Syngeneic NK-92 lines with constitutive expression of different GPCRs were harvested, resuspended in RPMI containing 1% FBS (NK migration media), and counted using Countess. 7.5 x 10^4^ cells in 75 μl were placed in each 8μm transwell insert (Corning HTS transwell 96-well plate, 3374). 200 μl NK migration media with varying concentrations of relevant ligands, as indicated in the figures and figure legends, was added to the bottom plate and incubated at 37 °C and 5% CO_2_ for 4 hours. For primary T and NK cell migration assay, a 3 μm transwell insert (Corning HTS transwell 96-well plate, 3385) was used instead of an 8 μm insert. The following ligands were used: Recombinant human IL-8 (Peprotech, 200-08M), Recombinant human C5a (Peprotech, 300- 70), 18:1 LysoPS (Avanti, 858143), 6-OAU (MedChem, HY-12764), and 7α,25-OHC (MedChem, HY-113962).

To test GPR183-dependent NK cell chemotaxis to cancer cell-derived factors, MDA-MB-231 cancer cell conditioned media was prepared as follows. 5 x 10^5^ MDA-MB-231 cells were seeded in a 6-well plate, treated with 40 ng/ml of human IFNγ (Peprotech, 300-02) and TNF (Peprotech, 300-01) in NK base media the following day, and incubated at 37 °C and 5% CO_2_ for additional 24 hours. MDA-MB-231 conditioned media was collected and centrifuged to remove cellular debris. 7.5 x 10^4^ NTC and GPR183^A+^ NK cells in 75 μl were placed in each 8μm transwell insert, and 200 μl MDA-MB-231 conditioned media was added to the bottom plate as a chemoattractant and incubated at 37 °C and 5% CO_2_ for 4 hours.

For competitive migration assay, NTC and GPR183^A+^ NK cells were stained with cytopainter orange (Abcam, ab176737) and deep red (Abcam, ab176736) dyes at room temperature for 20 min, respectively. After incubation, cells were washed with NK base media, counted, and then mixed 1:1. 7.5 x 10^4^ mixed cells in 75 μl were placed in each 8μm transwell insert (Corning HTS transwell 96-well plate, 3374). 200 μl NK migration media and 1:5 diluted tumor lysate were added to the bottom plate and incubated at 37 °C and 5% CO_2_ for 4 hours. 100 nM 7α,25-OHC was used as a positive control. For tumor lysate preparation, dissected tumor tissues were first weighed and submerged in PBS at 100 mg/ml. Then, tissues were ground using a tissue homogenizer (OMNI, TH115), and centrifuged at 4 °C for 10 min at 8000 rpm to eliminate cellular debris. The supernatant was collected, aliquoted, and cryopreserved at −80 °C until use.

In all transwell migration assays, quantification of migration was obtained as follows. After incubation, top inserts were removed, and migrated cells were collected and counted using a flow cytometer (NovoCyte, Quanteon) with cell counting beads (Biolegend, 424902). The migration index was determined by the fold change of each cell migrated to the chemoattractant compared to the medium. For competitive migration assay, the relative migration index was determined by (GPR183^tumorlysate or 7α,25-OHC^/Control^tumorlysate or 7α,25-OHC^)/ (GPR183^input^/Control^input^), where control are the NK or T cells transduced with a control construct (NTC sgRNA or Empty ORF).

### Quantification of in vivo NK infiltration

Seven to eight-week-old female NSG mice were subcutaneously injected with 5 × 10^6^ MDA-MB- 231 cells in a 1:1 mix of serum-free DMEM and Matrigel. A week after, when tumor size reached ∼50 mm^3^, tumor-bearing mice were randomized, and 5 × 10^6^ NK cells were adoptively transferred into tumor-bearing mice by intravenous tail-vein injection once a week for four weeks. Human IL- 2 (5000 U) was intraperitoneally administered daily for 4 consecutive days. Two days after the 4^th^ NK injection, tumors were dissected, minced into small pieces, digested with Collagenase type IV and DNase I for 45 min at 37 °C with agitation, and then filtered through a 100 μm cell strainer. Red blood cells were removed by using Red Blood Cell Lysing Buffer (Sigma, R7757) for 5 min at room temperature. The cell suspension was then diluted with 10 ml of PBS containing 2% FBS, passed through a 100 μm cell strainer, pelleted, and resuspended in NK base media. Single-cell suspension was further processed for flow cytometry staining to quantify tissue-infiltrated NK cells as described in the “Flow cytometry and cell sorting” section. 7-AAD^−^ mouse CD45^−^ GFP^+^ human CD45^+^ cells were identified as NK cells. Cell counting beads were used for precise cell counts.

### Cell proliferation assay

Control NTC (transduced with NTC sgRNAs) and GPR183^A+^ NK cells were stained with 1X Cytopainter deep red (Abcam, ab176736) at room temperature for 20 min. The cells were washed with NK base media, counted, and seeded at 10^5^ cells in 200 μl on 96 well plates. The level of cytopainter was monitored daily using a flow cytometer (NovoCyte, Quanteon) for 4 days. The relative mean fluorescent intensity was calculated.

### CD107a degranulation assay

5 × 10^5^ NTC and GPR183^A+^ NK cells were plated in NK base media supplemented with 1X Brefeldin A (BD GolgiPlug, 555029), 1X monensin (BD GolgiStop, 554724), anti-human CD107a-APC antibody (Biolegend, 328619, 1:200), and 1X cell activation cocktail (Biolegend, 423301). Four hours after activation, cells were washed with FACS buffer (PBS + 2% FBS), collected, and analyzed using a flow cytometer (NovoCyte, Quanteon).

### Intracellular IFNγ staining

5 × 10^5^ NTC and GPR183^A+^ NK cells were plated in NK base media supplemented with 1X Brefeldin A (BD GolgiPlug, 555029) and 1X cell activation cocktail (Biolegend, 423301). Four hours after activation, cells were washed with FACS buffer, fixed and permeabilized with Cytofix/Cytoperm kit (BD Cytofix/Cytoperm, 554714), and then stained with anti-human IFNγ- APC antibody (Biolegend, 506510, 1:200) at 4 °C for 20 min on ice. The cells were analyzed using a flow cytometer (NovoCyte, Quanteon).

### In vitro NK cell cytotoxicity assay and IFNγ-secretion quantification

MDA-MB-231 cells were seeded on a black-walled and clear-bottom 96 well plate (greiner, 655090). One day after cancer cell seeding, NK cells were added with varying effector-to-target ratios, as indicated in the figures and figure legends. After 48 hours of coculture, the supernatant was harvested and assessed with human IFNγ ELISA (Biolegend, 430104), following the manufacturer’s instruction. The remaining cells in the plate were washed with PBS five times, and the Prestoblue Cell Viability reagent (Thermo Fisher, A13262) was added to each well and incubated for 20 min. Fluorescent intensity was measured using a Tecan infinite M1000 plate reader.

### CAR construct design, cloning, and assessment of surface expression

Anti-hEpCAM CAR (**Figure 5c**) was cloned and used as follows. Lentiviral CAR backbone plasmid encoding anti-CD19-41bb-CD3ζ-EGFP genes was purchased from Addgene (135992). The original CD19-CAR construct was digested with FastDigest Bpil to replace anti-CD19-scFv- CD8hinge region with anti-hEpCAM-scFv-CD28hinge. The codon-optimized gene fragment encoding anti-hEpCAM-scFv-CD28 hinge was synthesized from Twist Bioscience and cloned into a digested backbone with Gibson assembly. High titer lentivirus of anti-hEpCAM CAR was generated and used for NK-92 transductions as described above. Transduction efficiency was measured by detecting GFP via flow cytometry. The surface expression of anti-hEpCAM CAR was assessed with Alexa Fluor 647-AffiniPure Goat Anti-Human IgG, F (ab’)_2_ Fragment Specific (Jackson Immuno Research Labs, 109605006, 1:100) via flow cytometry (**Supplementary Figure 9b**).

### In vivo disease outcome experiments

Seven to eight-week-old female NSG mice were subcutaneously injected with 5 × 10^6^ MDA-MB- 231 cells in a 1:1 mix of serum-free DMEM and Matrigel. A week after, when tumor size reached ∼50 mm^3^, tumor-bearing mice were randomized, and 10^7^ NK cells were adoptively transferred into tumor-bearing mice by intravenous tail-vein injection once a week for four weeks. Human IL-2 (5000 U) was intraperitoneally administered daily for 4 consecutive days. Tumor volumes were measured by mechanical caliper every 3 or 4 days and calculated with the following formula: V= 0.5 × length (L) × width^2^ (W).

### Flow cytometry and cell sorting

Cells were harvested and washed with FACS buffer (PBS + 2% FBS). Human TruStain FcX (Fc receptor blocker) (Biolegend, 422302, 1:50) was used to minimize nonspecific binding of antibodies. For live/dead cell stain, 7-AAD (Sigma-Aldrich, SML1633) was used. Cells were then stained for surface or intracellular markers at 4 °C in the dark for 20 min unless stated otherwise. For detecting surface levels of GPCRs, staining was performed at 37 °C in the dark for 30 min. The following antibodies were used: anti-human CD182 (CXCR2) (Biolegend, 320714), anti-human CD88 (C5AR1) (Biolegend, 344303), anti-human GPR183 (Biolegend, 368903), anti-human CD326 (EpCAM) (Biolegend, 324207), anti-human CD16 (Biolegend, 980104), anti-human CXCR6 (Biolegend, 356005), anti-human CD3 (Miltenyi Biotech, 130-113-134), anti-human CD8a (Miltenyi Biotech, 130-113-162), anti-human TCRα/β (Biolegend, 306717), anti-human CD56 (Miltenyi Biotech, 130-113-305), anti-human CD45 (Biolegend, 368510), and anti-mouse CD45 (Biolegend, 103116). Flow cytometry was performed on Quanteon, Penteon (NovoCyte), or Sony Biotechnology SH800S Cell Sorter. Cell sorting was performed using Sony Biotechnology SH800S Cell Sorter. All flow cytometry data were analyzed using FlowJo version 10.10.0.

### Quantitative PCR

Total RNA was isolated using Quick-RNA Microprep Kit (Zymo research, R1051) according to the manufacturer’s protocol. 1mg RNA was reverse transcribed to generate cDNA using QuantiTech Reverse Transcription kit (Qiagen, 205311). RT-qPCR was performed using PowerUp SYBR green master mix (Applied Biosystems, A25741) on a CFX96 or 384 Real-Time PCR system (Bio-Rad). Gene expression was normalized using the comparative threshold cycle method with human *GAPDH* as an endogenous control. All PCR primers used in this study are listed in **Supplementary Table 5**.

### CRISPR KO of EpCAM in MDA-MB-231

MDA-MB-231 cells stably expressing Cas9 were purchased from GeneCopoeia (SL515). Protospacer sequences of three NTC and three hEpCAM targeting sgRNAs were obtained from the Human CRISPR Knockout Pooled Library (GeCKO v2) and synthesized from IDT DNA. The oligos were annealed and cloned into the lentiGuide-Puro backbone (Addgene, 52963) and digested with FastDigest BsmBI (Thermo Scientific, FD0454). Equal amounts of the three hEpCAM sgRNA vectors were pooled and used for transfecting Lenti-X 293T cells to generate lentivirus. Cas9-expressing MDA-MB-231 cells were transduced with sgRNA and selected with 1 μg/ml puromycin. The surface expression of EpCAM was assessed to confirm gene knockout (**Supplementary Figure 9a,c**). The same procedure was repeated using the three NTC sgRNAs to generate control MDA-MB-231 cells. The oligo sequences are listed in **Supplementary Table 5**.

### Perturb-seq library and sequencing

Protospacer sequences of five NTC sgRNAs and three sgRNAs targeting top 8 hits from CRISPR activation screens were used to generate syngenetic NK-92 CRISPR/dCas9 cells with constitutive overexpression of top hits (**Supplementary Table 5**, as described in “Generating syngeneic NK- 92 lines with GPCR overexpression” section). The cells were pooled into a single cell suspension to include 20% NTC cells and 10% from each of the eight types of perturbed cells, according to the 10× ‘Single Cell Suspensions from Cultured Cell Lines for Single Cell RNA Sequencing’ protocol (10x Genomics, CG00054 Rev B) and processed on a 10x Chromium controller instrument. The libraries were prepared according to the Chromium Next GEM Single Cell 5′ Reagent Kits v2 (Dual Index) with Feature Barcode technology for CRISPR Screening protocol (10x Genomics, CG000510). Equimolar amounts of indexed libraries were pooled and sequenced on a NextSeq 2000 P3 in a paired-end run.

### Primary NK and CD8 T cell isolation, expansion, and engineering

Primary human NK and CD8 T cells were isolated from human whole blood buffy coats obtained from the Stanford Blood Center. First, PBMCs were purified using the Ficoll-Paque Premium Medium (Cytova, 17-5442-02) and Sepmate tubes (STEMCELL Technologies, 85450). For CD8 T cells, PBMCs were further processed with the EasySep Human CD8 T cell isolation kit (STEMCELL Technologies, 17953). Enriched CD8 T cells were cultured in RPMI-1640 with GlutaMax and HEPES medium (Gibco, 72400047) containing 10% heat-inactivated fetal bovine serum, 1% heat-inactivated human AB serum (MilliporeSigma, H4522), 5 mM of sodium pyruvate, 5 mM of non-essential amino acids, and 50 µM of b-mercaptoethanol (Sigma, M6250) at 1 × 10^6^ cells/mL in tissue-culture untreated 24-well plate. On day 0, T cells were activated with Dynabeads (Gibco, 11131D) at a 1:1 ratio for 3 days, followed by incubation with 100 U/ml IL-2. For T cell transduction, concentrated lentivirus was directly added to each well containing CD8 T cells and mixed by gentle pipetting on day 1 post Dynabeads activation. On day 3, beads were removed, and cells were split at 1 × 10^6^ cells/mL with fresh medium and IL-2. On day 5, T cells were harvested and sorted based on mCherry using Sony Biotechnology SH800S Cell Sorter. Sorted mCherry^+^ T cells were replated at 1 × 10^6^ cells/mL with fresh medium and IL-2 for 5 more days. For NK cells, PBMCs were further processed with the EasySep Human NK cell isolation kit (STEMCELL Technologies, 17955). Enriched NK cells were cultured in EL837 serum-free culture medium (EliteCell, ELM-1000) with EliteGro-Adv. (EliteCell, EPA-050), 5% heat-inactivated human AB serum (MilliporeSigma, H4522), 1000 U/ml hIL-2, and 100 ng/ml hIL-18 (biotech, 9124-IL) at 1 × 10^6^ cells/mL in tissue-culture untreated 24-well plate (day 0). On days 3 and 5, NK cells were replenished with fresh media containing IL-2 and IL-18. On day 7, concentrated lentivirus with 4 μg/ml polybrene (sigma, 107689-10G) was directly added to each well containing NK cells, followed by spin-infection at 30 °C for 90 min at 2500 rpm. Spin infection was repeated on day 8 to increase transduction efficiency. On day 10, NK cells were harvested and sorted based on mCherry using Sony Biotechnology SH800S Cell Sorter. Sorted mCherry^+^ NK cells were replated at 1 × 10^5^ cells/mL with fresh medium containing IL-2 and IL-18 for 3 days and used for in vitro migration assay.

### scRNA-Seq data analyses

ScRNA-Seq data previously generated from tumor and blood samples obtained from breast cancer patients was downloaded from the Gene Expression Omnibus (GEO; accession number GSE169246). For each cell type, differentially expressed genes (FDR < 0.01, log fold change > 0.1) were identified by comparing the tumor and blood samples using the Model-based Analysis of Single-cell Transcriptomics (MAST) test – a hurdle model tailored to scRNA-seq data. Cell types represented by less than 500 cells in either the tumor or blood samples were not considered in these analyses. The overall expression of the differentially expressed gene set shown in **Figure 1b** was computed with a normalization scheme to filter technical variation as previously described.

### CRISPR activation screen data analyses

Data from the CRISPR activation screens were analyzed using MAGeCK (version 0.5.9.4) (36,37). Fastq files were processed to align the reads to the sgRNA spacer sequences, generating a counts matrix of sgRNA by samples. Using MAGeCK, the variance of read counts was estimated by sharing information across the different sgRNAs and by using the NTC sgRNAs to adjust and control for technical variation across the samples. Effect size and *p*-values were computed based on a negative binomial statistical model for every target gene by testing whether sgRNA abundance differs significantly between two groups of samples. All *p*-values were corrected for multiple hypotheses testing using the BH correction, and FDRs are reported (**Supplementary Table 2** and **3**). In the in vivo screens, NK cells in the tumor were compared to NK cells in the lung and to the reference NK pool. The data from each of the three in vivo screens was analyzed separately and subsequently combined via Fisher statistic *p*-values as the summary statistics. The results are provided both separately and when combined to summary statistics (**Supplementary Table 2**). In the in vitro chemotaxis screens, NK cells collected from the bottom insert of the transwell (4 hours after placement at the top insert) were compared to the initial NK cell pool (cultured for 4 hours). The screen was conducted twice, once with MDA-MD-231 supernatant as the chemoattractant and once with MCF7 supernatant as the chemoattractant. Fisher statistic *p*- values were computed as the summary statistics combining the results from the two types of chemotaxis screens. The results are provided both separately and when combined to summary statistics (**Supplementary Table 3**). In the spheroid screens, NK cells residing within MCF7 spheroids after 6, 24, and 48 hours were compared to NK cells cultured for 6, 24, and 48 hours, respectively. Fisher statistic *p*-values were computed as the summary statistics to combine the results from all three time points. The results are provided both separately and when combined to summary statistics (**Supplementary Table 3**).

### Perturb-seq data analyses

Raw fastq files were processed using the cellranger pipeline (10x Genomics Cell Ranger 7.1.0) to align reads to the human genome (GRCh38) and to the sgRNA spacer sequences to generate two counts matrixes, one of gene expression (genes by cells) and another of sgRNA detection (sgRNA by cells). Gene expression counts were converted to transcript per million (TPM) values and log transformed. Cells were assigned to sgRNAs using the default cutoffs, and only cells with a single confidently detected sgRNA were used in all downstream analyses. Seurat R package was used to cluster the TPM gene expression profiles using the Shared Nearest Neighbor (SNN) clustering algorithm (Waltman and van Eck, 2013). For each of the eight genes targeted in the screen, a MAST analysis was performed to identify differentially expressed genes in the cells carrying one of the target sgRNAs in comparison to the control cells (i.e., cells carrying a NTC sgRNA absolute log fold-change > 0.1 and BH FDR < 0.01). The z-score vector for each target was defined as |log_10_ *p*| * *sign* (*logFC*) and used as the gene ranking for gene set enrichment analyses (GSEA), using the fast GSEA package (*fgsea,* version 1.24.0).

## Data availability

The data collected in this study will be made publicly available upon manuscript publication through GEO, Zenodo, and the Single Cell Portal. Data deposited on the Single Cell Portal will also be available for interactive web visualization.

## Code availability

Code to reproduce the results presented in this study will be made available as a GitHub repository at https://github.com/Jerby-Lab.

